# Characterisation of HIF-dependent alternative isoforms in pancreatic cancer

**DOI:** 10.1101/826156

**Authors:** Philipp Markolin, Natalie Davidson, Christian K. Hirt, Christophe D. Chabbert, Nicola Zamboni, Gerald Schwank, Wilhelm Krek, Gunnar Rätsch

**Author notes:** The authors wish it to be known that, in their opinion, the first 2 authors should be regarded as joint First Authors. Deceased (Wilhelm Krek). Christophe D. Chabbert, Roche Innovation Center Zürich, Wagistrasse 10, 8952 Schlieren, Switzerland Gerald Schwank, Universität Zürich Institut für Pharmakologie und Toxikologie, Winterthurerstrasse 190, Y17 J34, CH-8057 Zürich.

## Abstract

Intra-tumor hypoxia is a common feature in many solid cancers. Although transcriptional targets of hypoxia-inducible factors (HIFs) have been well characterized, alternative splicing or processing of pre-mRNA transcripts which occurs during hypoxia and subsequent HIF stabilization is much less understood. Here, we identify HIF-dependent alternative splicing events after whole transcriptome sequencing in pancreatic cancer cells exposed to hypoxia with and without downregulation of the aryl hydrocarbon receptor nuclear translocator (ARNT), a protein required for HIFs to form a transcriptionally active dimer. We correlate the discovered hypoxia-driven events with available sequencing data from pan-cancer TCGA patient cohorts to select a narrow set of putative biologically relevant splice events for experimental validation. We validate a small set of candidate HIF-dependent alternative splicing events in multiple human cancer cell lines as well as patient-derived human pancreatic cancer organoids. Lastly, we report the discovery of a HIF-dependent mechanism to produce a hypoxia-dependent, long and coding isoform of the UDP-N-acetylglucosamine transporter SLC35A3.

## Introduction

Oxygen is essential for eukaryotic life. Throughout the evolutionary history of life, it was necessary for organisms to evolve and integrate elaborate oxygen sensing and adaptation systems into cellular pathways to ensure cellular survival during periods of reduced oxygen availability, hypoxia [1,2]. The most important and well-studied proteins involved in oxygen sensing and signaling include hypoxia-inducible factors (HIFs) and their regulators [3]. HIF1α and HIF2α proteins are a class of transcriptional activators which, under hypoxic conditions, dimerize with ARNT/HIF1β [3,4] and form a transactivation complex with p300/CBP [5] to regulate gene expression of thousands of genes involved in various cellular pathways, including angiogenesis [6,7], proliferation [7–9], metabolism [10–12], apoptosis [13,14] and DNA repair [15].

The versatility of HIF signaling also plays an important role in cancer biology [16–18]. Many solid tumors experience stages of intermediate hypoxia during neoplastic growth with subsequent upregulation of the HIF pathway [18–20]. Furthermore, it has been repeatedly shown that cancer cells, utilizing HIF transcriptional programs, are able to gain competitive advantages over normal cells through metabolic adaptation, growth optimization, immune system evasion, and better survival in pathophysiological microenvironments [20–25].

More recently, hypoxia has been reported to affect alternative splicing of HIF and non-HIF target genes [26,27], and that HIF activity but not hypoxia *per se* is necessary and sufficient to regulate RNA splicing of hypoxia inducible genes [26].

Defective or aberrant alternative splicing is a prominent and widespread feature in cancer biology [28,29]. Alternative splicing is abundant in higher eukaryotes, greatly expanding the diversity of the human transcriptome and proteome [29,30]. It has been estimated that over 95% of human genes undergo alternative splicing [31] and that up to 50% of disease-causing mutations influence splicing [32,33]. Consequently, it was reported that pathologically altered alternatively spliced genes are involved in almost every aspect of cancer biology, including proliferation, differentiation, cell cycle control, apoptosis, motility, invasion, angiogenesis, and metabolism [29,33–36].

While genomic data resources are now available for many cancer types, broad studies investigating environmental context-driven alternative splicing or RNA processing in various cancer types are scarce [34,37].

Given the crucial role of HIFs to implement adaptation to hypoxia, we set out to perform whole transcriptome sequencing in pancreatic cancer cells with and without perturbation of HIF transcriptional activity in normoxic (21% O2) and hypoxic (1% O2) settings. We analyze transcriptome data using SplAdder [38] and compare identified alternative splicing events to TCGA patient data. To identify biologically relevant HIF-driven splice events, we performed an association study to identify which of our events are significantly associated with the hypoxic state of the patient. Next, we experimentally validate hypoxia-induced and HIF-dependent splicing events in multiple *in vitro* systems, including clinically relevant patient-derived human pancreatic cancer organoids. We report the discovery of a HIF-dependent mechanism to produce a long, coding isoform of the UDP-N-Acetylglucosamine (UDP-GluNAc) transporter SLC35A3 (SLC35A3-L). Lastly, we investigate metabolic adaptations to siRNA mediated selective knockdown of SLC35A3-L in both normoxia and hypoxia.

## Methods

### RNA preparation and sequencing

AsPC-1 cells were seeded in 6-well plates in triplicates and transfected with siScramble or siARNT using Lipofectamine RNAiMAX. The next day, cells were split 1:2 and transferred to a hypoxic chamber (Baker Ruskinn SCI-tive) at 1% O_2_ for 24h or kept under normoxic culture conditions. 48h after transfection, cells were harvested using the Nucleospin® RNA extraction protocol (Machery-Nagel). An aliquot was taken to measure RNA concentration and perform quality control, the rest was immediately frozen at −80°C until shipping to the Sequencing Facility (Genomics Facility Basel).

RNA concentration and quality was assessed with Ribogreen and QC length profiling by the facility before library generation (TruSeq® Illumina mRNA (Poly(A) enrichment) stranded). Sequencing was performed on three lanes (HiSeq, SR125, 250-300 M reads per lane) and read sequences stored in FASTQ format.

### Data processing

Sequencing reads were trimmed using trimmomatic [39] to remove residual Illumina oligonucleotide sequences on the 3’ end of each tag (options: ILLUMINACLIP: TruSeq3-SE.fa:2:30:8 LEADING:25 TRAILING:3 SLIDINGWINDOW:4:15 MINLEN:36). The remaining reads were aligned to the GRCh38.p10 human genome assembly using STAR 2.4.2a [38,40] together with the Genecode v26 gene annotation (default parameters). The search for novel junctions was allowed during the mapping step. Gene level read counts were generated using the QoRTs software v1.2.42 [39,41] after excluding reads with multiple alignments (MAPK score less than 255).

### Differential gene expression analysis

Gene count tables were loaded into R as a DESeq2 [42] object to conduct differential gene expression analysis. Genes with very low counts (less than 20 mapped reads across all 12 samples) were excluded and we used the subsequently estimated size factors to correct for differences in library size. Following the standard DESeq2 workflow [42,43], changes in gene expression were modelled using one variable accounting for differences in genotype (wild type or ARNT knock down) and oxygen levels (normoxia or hypoxia) in each group of samples. Genes whose mRNA expression was impacted by ARNT in hypoxic and normoxic conditions respectively were identified when comparing each relevant group of samples (WT, normoxic conditions and WT, hypoxic conditions for example). Conversely, genes with an ARNT-dependent expression under hypoxic conditions were identified by comparing the log2 fold changes during hypoxic responses for both genotypes. For each contrast of interest, results were extracted with the DESeq2 results function and multiple testing correction performed using the Benjamini Hochberg procedure [44]. Only genes with an adjusted p-value lower than 0.05 and an absolute log2 fold change of at least 0.58 (1.5-fold increase or decrease) were considered differentially expressed between conditions.

### Gene Ontology enrichment

In order to account for potential biases in gene ontology enrichment analysis, the matchIt function [45] was used to generate a gene set with expression and width distributions identical to that of the differentially expressed genes. This set constituted a background for the enrichment estimations performed using the weight algorithm from the topGO package [46]. The significance of each enrichment was assessed using a Fisher statistic and GO terms were then ordered by p-values within each ontology independently (molecular functions, cellular compartments and biological processes).

### SplAdder/Differential Splicing

To identify all splice events on the aligned bam files, SplAdder [38] was used with gencode annotation version 19. Splice events considered were: exon skips, alternative 3’/5’, and intron retentions. We considered only events with highest confidence, as defined by SplAdder. All parameters used are shown here: -M merge_graphs -t exon_skip, intron_retention, alt_3prime, alt_5prime, mult_exon_skip, mutex_exons -c 3.

To identify which events are HIF-dependent, we first identify which junctions represent a specific splice event, the junction including or excluding the alternative exon part. Junction identification for each event was done by SplAdder’s differential junction count test. The junction chosen to represent the event was the junction that had the most significant difference between the +/-HIF conditions as reported by SplAdder’s test. Once the junction of interest was identified for each splice event, DESeq2 [42] was used to estimate the junction count dispersions. glm.nb [47] from the MASS R package was then used to test for HIF dependence, independent of expression between conditions. The experiment was modeled as:

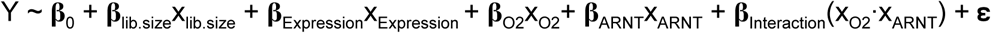

Y is the raw junction read counts, x_Expression_ is the library-size corrected expression of the gene where the junction is located, x_O2_ denotes whether the sample is exposed to hypoxic or normoxic conditions, x_ARNT_ denoted whether the sample had siARNT or siCTRL added. All **β** terms are the associated coefficients in the model for each feature. The significance of HIF-dependence is defined as the Benjamini-Hochberg corrected significance of the interaction term, **β**_Interaction_, in our model. Furthermore, to ensure the events were not driven by expression changes, we additionally filtered events that had a HIF-dependent change in expression with an adjusted p-value ≤ 0.05 and log2FC > 1.

To identify events that had HIF-dependent junction and mRNA count changes, we modeled our experiment in DESeq2 as:

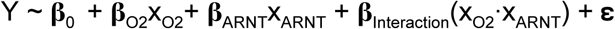

Y is the raw junction and expression counts, and all other variables are the same as defined in the previous model. Again, the significance of HIF-dependence is defined as the Benjamini-Hochberg corrected significance of the interaction term, **β**_Interaction_, in our model.

Only splicing events that had an FDR corrected p-value < 0.1 and a ΔPSI (percent spliced in) > 0.05 were selected for analysis in the TCGA cohort. ΔPSI is defined as:

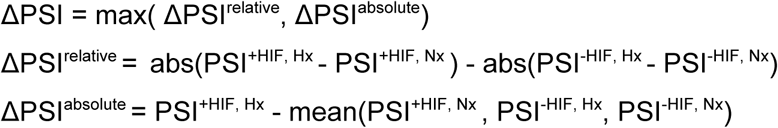

### Correlating Splicing Events with Hypoxia Score

To identify the hypoxia score across all TCGA samples [48], we used the R package ssPATHS[49]. We used the hypoxic genes defined in the function get_hypoxia_genes to calculate the per-patient hypoxia score. Using the hypoxia score across all samples in TCGA with available RNA-Seq measurements, we associated the hypoxia score with the PSI or expression values of the events we identified as HIF-specific in our experiment using the two models displayed below. All expression and PSI values were taken from SplAdder.

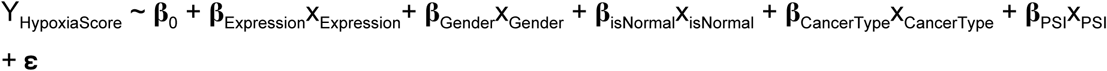

where **ε** ∼ Gaussian(*μ,σ*^2^)

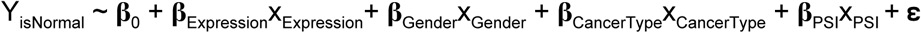

where **ε** ∼ Binomial(n, p)

Only events that were significant in both tests were displayed in Figure 2d and Supplementary Figure S2. For all events, excluding SLC35A3, x_PSI_ is the PSI of the event. For SLC35A3, we used the expression of the first exon as x_PSI_. This was done because the expression of the first three exons is low in most samples, which leads to very few junction read counts and unstable regression estimates. To avoid this, we only consider the expression of the first exon, which is still a measure for the presence of the long isoform of interest. x_Expression_ is the library-size normalized expression of the gene, x_Gender_ is the gender of the patient, x_isNormal_ denotes if the sample is an adjacent normal or a tumor sample, x_CancerType_ is the TCGA study abbreviation assigned to the sample, and x_PSI_ is the PSI for the event. The significance of the association is defined as the the Benjamini-Hochberg corrected significance **β**_PSI_, in our model.

**Figure 1:**
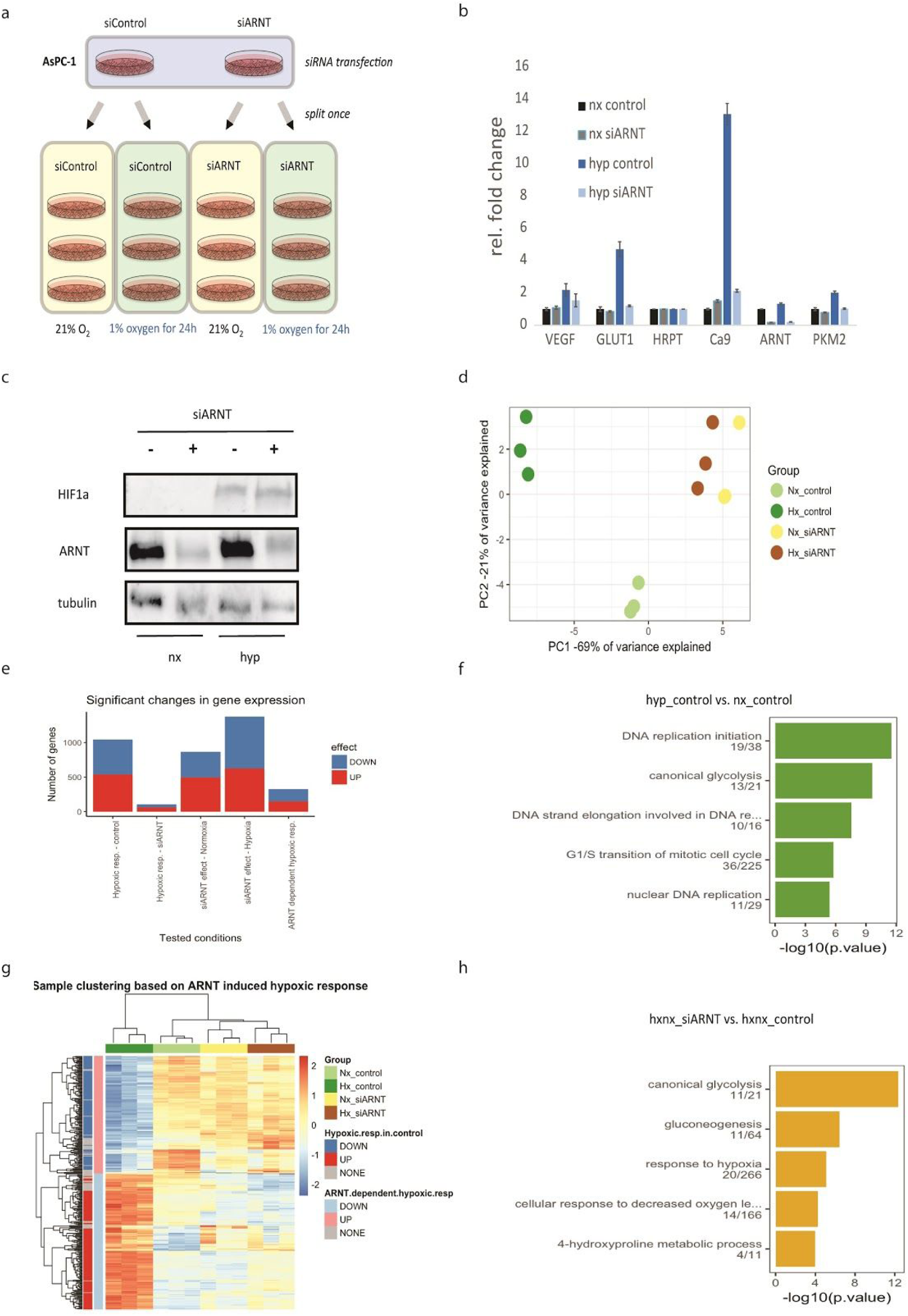
Global gene expression changes during hypoxia. **a.** Schematic of experimental setup for RNA seq. **b.** qRT-PCR validation of HIF-target genes and ARNT knockdown. **c.** Western blot validation of HIF protein expression and ARNT knockdown of sequenced samples **d.** Principal component analysis of sample similarity in 2D projection. **e.** Number of significant gene expression changes between conditions. **f.** Gene ontology of top 5 enriched pathways during hypoxia. **g.** Hierarchical clustered heat map of significantly changed HIF-dependent genes over all comparison conditions. **h**. Gene ontology of top 5 enriched pathways for hypoxia inducible and ARNT dependent genes. Nx, normoxia; Hyp, hypoxia; siARNT; siRNA mediated knockdown of aryl-receptor nuclear translocator (ARNT); siControl, siRNA control (nontargeting)

**Figure 2.**
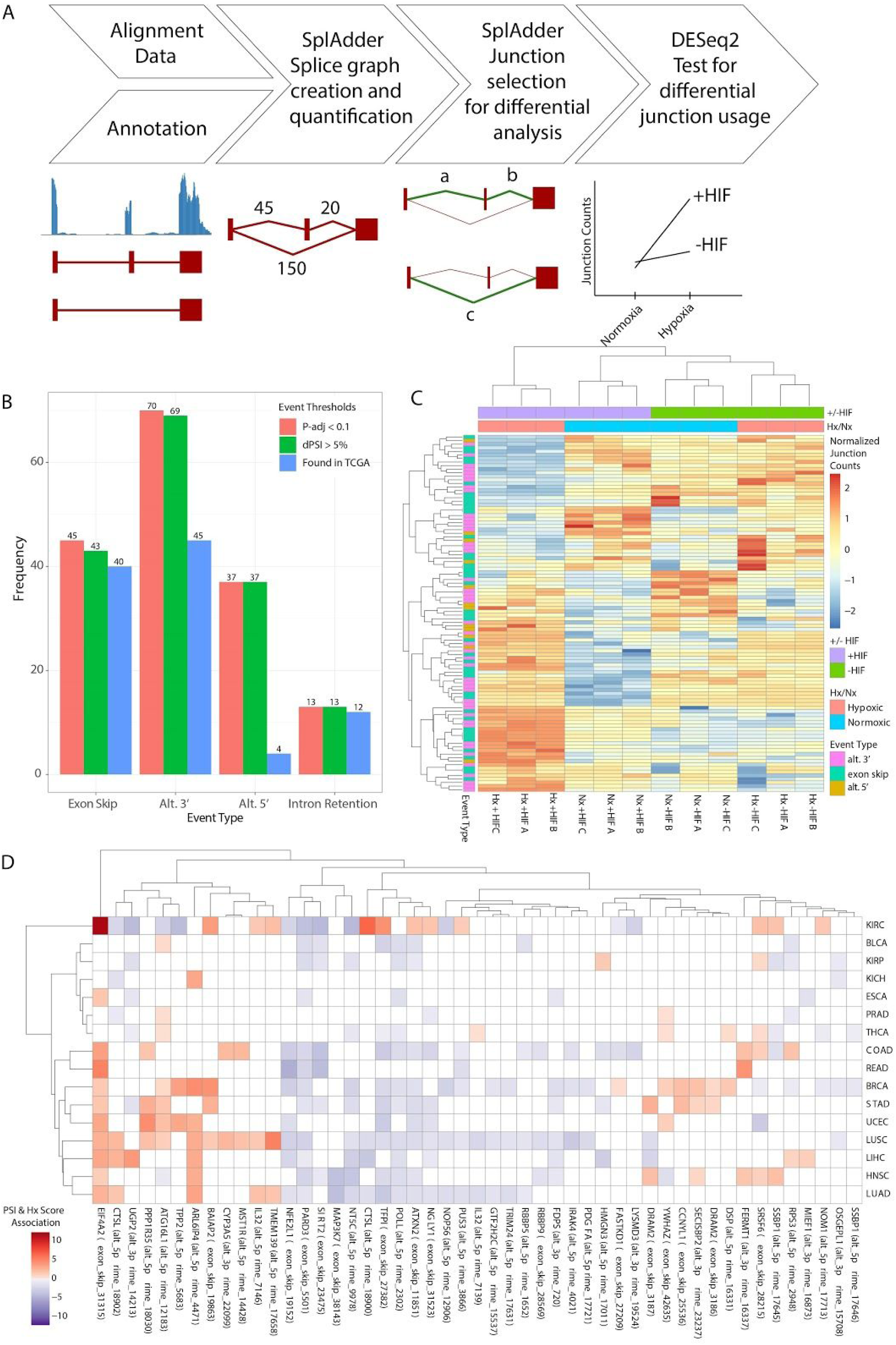
Identification of hypoxia-inducible alternative splicing events. **a.** SplAdder pipeline. **b.** Number of alternative splicing events identified using p-adjusted (pink) alone or p-adjusted and PSI >5% (green) combined as threshold. Number of remaining events after adding the additional filter that the events are observed in TCGA are depicted in blue. **c.** Heat map of splice junction read counts centered and scaled across conditions for identified alternative splicing events after hierarchical clustering. We observe that the triplicate cluster with one another in each condition showing that we have minimal within-condition variance. Furthermore, we find that the HIF-activated samples cluster away from all other samples, indicating that our significant events identify a HIF-specific behavior. **d.** Heatmap of association results between PSI of our events of interest and an estimated hypoxia score in the TCGA cohort. We find that our events are significantly associated in multiple cancer types, with some events (EIF4A2 and PARD3) maintaining a constant direction of association across all cancer types.

### RNA harvest & isolation

Cells were washed 2x with cold PBS and subsequently scrapped in 350 µl lysis buffer containing 10 mM Tris(2-carboxyethyl)phosphine (TCEP) according to Nucleospin® RNA extraction protocol (Machery-Nagel). In short, the lysates were filtrated to reduce viscosity, then 350 µl ethanol (70%) was added and the suspension was mixed. The suspension was placed in a RNA column and centrifuged at 11.000 × g for 30 seconds. The membrane-bound RNA was desalted with 350 µl MDB buffer and DNA was digested with rDNase at RT for 15-20 minutes. Finally, the RNA was washed in 3 washing steps and eluted in 50 µl RNase-free H_2_0 and subsequently used for cDNA synthesis or frozen at −20 °C. RNA concentrations were measured using Nanodrop 1000 Spectrophotometer (Thermo Scientific).

### cDNA synthesis and quantitative real-time PCR

The RNA was reverse transcribed using high-capacity cDNA reverse transcription protocol (Applied Biosystems™). 1 µg total RNA was diluted in 10 µl nuclease-free H_2_O for each sample and mixed with 2x reverse transcriptase master mix including dNTPS, RT random primers and RNase inhibitor. The mix was incubated in a thermal cycler for 10 min at 25°C followed by 120 min at 37°C and inactivated by 85°C for 5 minutes. The resulting cDNA was diluted 1:10 with ddH_2_O stored at 4 °C.

Quantitative real-time PCR (qRT-PCR) reactions were prepared using Roche Lightcycler 480 SYBR Green® Master (Roche) as recommended by the manufacturer. PCR was performed on a Lightcycler 480 machine (Roche) and Ct values were normalized to the housekeeping gene HPRT (hypoxanthine-guanine phosphoribosyl transferase) [50]. All qRT-primers were either self-designed or sequences were taken from primerbank website (Harvard) and ordered by Microsynth. Primer sequences used for this study can be found in the appendix (Primer list).

### Immunofluorescent stainings

Immunofluorescent stainings were performed as described previously. HUH7 monolayers were fixated with 4% Paraformaldehyde (PFA) in PBS for 10 minutes and washed with PBS 3x afterwards. The reaction was stopped with 0.1M Glycine in PBS incubated for 10 minutes and washed again 3x with PBS. The cells were then permeabilized with 0.2% TX-100 in PBS for 10 minutes at room temperature and washed again after with PBS 3x. For incubation with primary antibody (diluted in 1:100 in PBS + 0.05 % Tween) a humid chamber on parafilm upside down for 1 hour at RT. Cells were washed 3 x with PBS before incubation with secondary antibody and DAPI (diluted in 1:1000 in PBS + 0.05 % Tween) in a humid chamber on parafilm upside down for 1 hour at RT in the dark. Finally, cells were washed 3x with PBS and mounted on a coverslip with a drop of Mowiol mounting medium and sealed with nail polish to prevent drying and movement under the microscope.

### siRNA transfection with Lipofectamine RNAiMAX

A transfection master mix was prepared using 250 uL Opti-MEM + 5 uL RNAiMAX per reaction and incubated for 5 min at RT. In parallel, a siRNA mix was prepared using 250 uL Opti-MEM + 5 uL siRNA (20mM stock) per reaction. Both mixes were combined after 5 min and incubated for another 20 min at RT. In the meantime, cells were trypsinized, resuspended in DMEM and counted.

500 µl of reaction mix per 1 well of a 6-well plate was added and then 2.5 * 10^5 cells were filled on top, diluted to yield a total volume of 2.5 ml per well. On the next day, cells were split 1:2 using 250 µl trypsin per well and incubated for 24-48 h before harvest.

### Lipofectamine 2000 transfections

Cells were seeded a day before transfection to reach a high confluency (95%). Depending on cell line, 1-3 µg plasmid DNA diluted in 250 µl optiMEM were prepared per 2.5 * 10^5 cells in a ratio of 1:2 or 1:3 with Lipofectamine 2000 (1mg/ml). Both DNA and Lipofectamine 2000 dilutions were mixed and incubated for at least 30 min (longer for bigger plasmids) before adding to cells. The reagents were mixed by swirling. After 4 h of incubation, the medium was changed to complete DMEM. Cells were harvested at least 24h after transfection.

### PDAC Organoid Line Generation

Human PDAC organoids were isolated as previously described [48,51]. Pathological tissue specimens from the tumor mass after surgical resection was placed in complete pancreas medium containing 5mg/ml Collagenase II. Tissue was digested at 37°C while shaking for 5-12h. Subsequently the digestion was blocked with cold Advanced DMEM supplemented with 10mM Hepes, 1x Glutamax, 1x Penicillin/Streptomycin, centrifuged at 120g and plated in 20ul Matrigel drops. After gelling pancreas growth medium containing RhoKinase inhibitor was added.

### Organoid Culture

Human PDAC organoid lines were cultured in AdDMEM/F12 medium (Life Technologies) supplemented with 10mM Hepes, 1x Glutamax, 1% Penicillin/Streptomycin, 1x B27 without vitamin A (Life Technologies), 1.25mM n-Acetyl-Cysteine (Sigma), 10uM Nicotinamide (Sigma), 50ng/ml EGF (Peprotech), 100ng/ml FGF1 (Peprotech), 10nM Gastrin (Tocris), 0.5 μM A83.01 (Tocris), 1 μM PGE2 (Peprotech), 50% Wnt3a (conditioned medium), 10% Noggin (conditioned medium), 10% R-spondin (conditioned medium) and 100 μg/ml Primocin (InvivoGen).

Organoids were split approximately once per week either by trypsinisation or by mechanical dissociation, followed by 5 minutes centrifugation at 120 rcf at 4°C. The cell pellet was then resuspended matrigel.

### Protein harvest

Protein lysates from cell or organoid culture were extracted using RIPA lysis protocol (Bethyl Laboratories). Before use, Phosphostop® & Complete protease inhibitor® cocktail mix (Roche) and 1 mM DTT was added to RIPA lysis buffer. Cells were grown to approximate 80-90% confluence in tissue culture plates. The culture medium was aspirated carefully and the monolayers were washed 2x with ice cold PBS. The cells were scrapped in 1 ml of PBS and centrifuged at 4000 rpm for 3 minutes, then the supernatant was aspirated and 1 ml RIPA lysis buffer (for 10 cm culture dishes) was added to the cell pellet and resuspended by pipetting up and down several times. The cell-lysis mix was incubated for 30 min at 4 °C on ice and then centrifuged at 4 °C for 15 min at maximum speed to pellet debris. The supernatant was extracted and transferred to a fresh Eppendorf tube and 2 µl were taken to measure total protein amount using Bradford assay. The samples were stored at −20 °C (short term) or −80 °C (long term).

### Western Blotting

10-40 µg of total protein lysate were separated by SDS-PAGE on 8% or 10% polyacrylamide minigels (BioRad) and transferred onto a nitrocellulose membrane (GE healthcare) by semi-dry transfer as described previously [52]. Transfer efficiency was checked with reversible Ponceau S (BioRad) staining. After removal of Ponceau S with 1x TBST (10 mM Tris base, 0.9 % w/v NaCl, 0.1 % Tween-20) the membranes were incubated for 2 h at RT in 5 % milk (in PBS) on a shaking platform.

Primary antibodies were incubated overnight at 4 °C on a rolling platform (∼ 20 rpm). The next day, antibody solution was retrieved (reuseable up to 5 times) and the membranes were washed 3 times for 10 minutes in 1x TBST. Subsequently, the membranes were incubated with HRP-coupled secondary antibodies (Invitrogen) for the respective species (1:5000 for anti-mouse and anti-rabbit in 5% milk/TBST) for at least 1 hour at room temperature on a rolling platform. After incubation, the membranes were washed again 3 times for 10 minutes with 1x TBST and proteins were visualized using SuperSignal West Pico Chemiluminescent Substrate (Thermo Scientific). Signals were digitally captured on a Fusion Solo S (Witec) machine.

The following antibodies were used: mouse monoclonal ARNT1 (cat. No. 611079, BD Biosciences), rabbit polyclonal HIF1α (cat. No. NB-100-47, Novus Biologicals), rabbit polyclonal GLUT1 (cat. No. 07-1401, Millipore), mouse monoclonal y-Tubulin (cat. No. T6557, Sigma), mouse monoclonal SF3B1 (cat. No. D221-3, MBL international), rabbit monoclonal SRSF1 (cat. No. 5764-1, Epitomics), rabbit polyclonal SRSF7 (cat. No. sc-28722, Santa Cruz), goat polyclonal hnRNP H (cat. No. sc-10042, Santa Cruz), mouse monoclonal SLC35A3 (cat. No. WH002344M1, Merck).

### Metabolomics

Cells were transfected with siRNA using RNAiMAX protocol (quadruplets). On the next day, cells were trypsinized and equally split 1:2 in separate 6-well plates for normoxia and hypoxia. After 60 min of waiting for cells to reattach, the corresponding 6-well plates were transferred to a hypoxic chamber and exposed to 24 h of 1 % O_2_ or maintained in normoxia at 21 % O_2_.

After incubation, cells were washed 2x quickly with ammonium carbonate solution (Sigma #207861) (pre-warmed at 37°C) and aspirated completely. Next, the bottom of the 6-well plates was dipped into liquid nitrogen for 60 sec to snapfreeze the cells. Subsequently, the cells were immediately processed to extract metabolites with 400 µl pre-cooled 40:40:20 acetonitril:methanol:water mix at −20°C for 10 min. The extraction solution was then collected in separate Eppendorf tubes and a second extraction with 400 µl, −20°C, 10 min was performed. The extraction solution was once again transferred to the Eppendorf tubes and then stored at −80°C until use.

## Results

### HIF proteins are essential to implement hypoxia-induced gene expression programs in PDAC cells

To assess the contribution of all three HIFα proteins to gene expression and alternative splicing output during hypoxia, we knocked down their dimerization partner ARNT with ARNT-specific siRNA in the human pancreatic cancer cell line AsPC1. Furthermore, we exposed the cell lines to normoxic (21% O_2_, 24h) or chronic hypoxic (1% O_2_, 24h) conditions (Fig. 1a). We verified >90% siRNA-mediated knockdown of ARNT in both RNA transcript and protein levels (Fig. 1b and 1c). Furthermore, we assessed the expression of canonical HIF target genes VEGF, GLUT1, CAIX and PKM2 (Fig. 1b). As expected, all tested HIF target genes increased when exposed to hypoxia but failed to increase in hypoxia when ARNT protein levels were abolished. Finally, we verified that HIF1α protein is still stabilized in hypoxia under ARNT depletion (Fig. 1c). Taken together, we concluded that ARNT knockdown in AsPC-1 cells is sufficient to impair transcriptional adaptation to hypoxia even when HIFα proteins are stabilized.

Next, stranded libraries were constructed from the RNA extracted from each sample and sequenced. On average, >60 million 125bp single-end reads per sample were mapped to the human genome (hg19) for gene expression and alternative splicing analysis. We performed a principal component analysis (Fig. 1d) on read counts over all samples and found that 90% of variance can be explained by the first 2 components.

Subsequently, we assessed differences in gene expression (up- and downregulated) by pairwise comparisons between the groups (Fig. 1e). We observe a strong hypoxia-response (adjusted p-value < 0.05, log2 fold change > 0.58, see Methods) in 1043 genes (539 up, 504 down) being differentially expressed when comparing normoxia control to hypoxia control (Fig 1e, left panel). In comparison, normoxia siARNT vs. hypoxia siARNT only altered expression of 106 genes (61 up, 45 down). Additionally, we also observed a strong effect of the siRNA knock down with significant changes (siControl vs. siARNT) in 864 genes (494 up, 370 down) in normoxia and (1375 genes; 627 up, 748 down) in hypoxia.

These changes were expected since ARNT is also prominently involved in aryl receptor signaling [53] and has been reported to regulate enzymes in xenobiotic metabolism [54]. Finally, we stratify a subset of 325 genes (174 up, 151 down) where expression changed in a ‘HIF-controlled’ manner characterized by both hypoxia-dependency and ARNT susceptibility (Fig 1e, right panel).

Next, we performed gene ontology and pathway analysis for baseline hypoxia and HIF-controlled gene subsets, respectively (Fig. 1f and 1g). Differentially expressed genes found by comparing normoxia control *vs* hypoxia control showed an enrichment in genes involved in DNA replication processes and G1/2 transition, as well as canonical glycolysis (Fig 1f). This pathway enrichment is consistent with reports that hypoxia arrests cell cycle [8,55,56] and reprograms glycolysis [10,12,57].

Notably, when we narrowed our analysis to the ‘HIF-controlled’ set of 325 genes which are ARNT-dependent and hypoxia-inducible, we found that the top five enriched pathways (canonical glycolysis, gluconeogenesis, response to hypoxia, cellular response to decreased oxygen and 4-hydroxyproline metabolism) are all involved in metabolic processes. This underscores the role of HIF proteins as metabolic master regulators during cellular hypoxia in pancreatic cancer cells.

### Identification of hypoxia-induced alternative splicing events

Next, we set out to find alternative splicing events between experimental conditions in our transcriptome data. We used SplAdder [38] after read alignment (STAR 2.4.2a, see Methods) to create an augmented splicing graph from which we extracted splicing events. The extracted splicing events were further quantified and splice-junction reads were selected for differential analysis between the samples. Finally, DESeq2 [42] and glm.nb [58] was used to test for HIF-dependent junction usage, independent of expression changes. (Fig 2a).

Using junction read counts, we identified 165 significant alternative splicing events with an adjusted p-value < 0.10 (see Methods). Alternative 3’usage (70 events) and exon skipping (45 events) and were the most prominent splicing changes followed by alternative 5’usage (37 events), intron retention (13 events) (Fig 2b, red). To ensure a greater biological relevance of retained events, we only considered events with a change in percentage-spliced-in (PSI) >5% (Fig. 2b, green). Further details on our PSI thresholding are given in the Methods section. Lastly, as we were interested in alternative isoforms found in human cancer patients, we filtered by retaining events that were observed in at least one patient in The Cancer Genome Atlas (TCGA). Notably, while 40/43 of exon skips and 12/13 intron retentions identified by Spladder are recapitulated in TCGA cancer cohorts, we could only find evidence for 45/69 alternative 3’ events and 4/37 alternative 5’ events (Fig 2b, blue).

We performed hierarchical clustering on the junction read counts of the events which we identified as HIF-specific (Fig 2c) and we observed a clustering for the biological triplicates over all experimental conditions. We also find a clear cluster separation between hypoxic samples (lanes 1-3) and normoxic or ARNT-perturbed samples (lanes 4-12), underscoring the crucial role of HIFs’ in implementing transcriptome changes during hypoxia.

Our next goal was to identify which of our events were putative cancer relevant HIF-driven splice events. To identify these we performed two more filtering steps. Firstly, again taking advantage of TCGA, we identified which of our events are significantly associated with the hypoxic state of the patients. Utilizing TCGA expression data, we used the R package ssPATHS [49] to estimate a per sample hypoxia score. We then tested the association between the PSI of our identified HIF-dependent events and the samples hypoxia score across multiple tumor backgrounds (Fig. 2d). Secondly, we manually identified which HIF-dependent alternative splicing events had a high chance of affecting biological functions by generating alternative transcripts (Supplementary table T1).

Furthermore, we assessed promising candidate events in human pancreatic cancer organoids subjected to hypoxia. We used human PDAC organoids derived from pancreatic cancer patients and 3D cultured in a matrigel matrix. Organoids form spherical structures (Supplementary Fig. S1) reportedly preserve many features of PDAC which cannot be reliably recapitulated in 2D culture systems [59]. Since alternative splicing cannot be compared between mouse and human, PDAC organoid models provide a good compromise between experimental and clinical applications [60,61]. For our experiment, we used 3 different PDAC organoids lines (PaCa-4/-6/-8) derived from genetically distinct PDAC patients and exposed them to hypoxia (1% O_2_, 24h). We designed a set of junction and exon-specific primers (Fig. 3a) to validate several exon skipping events identified by SplAdder.

**Figure 3.**
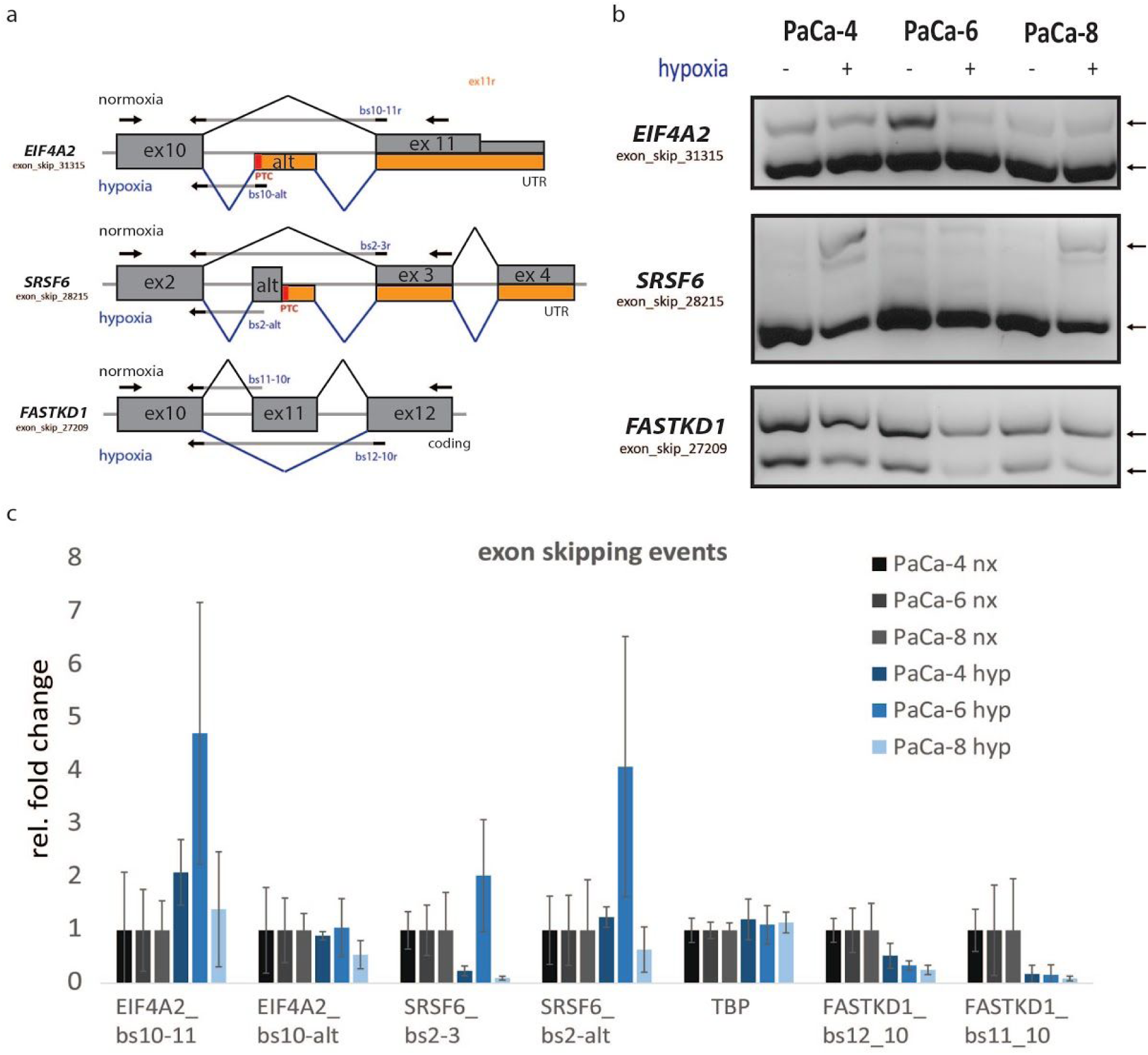
Validation of hypoxia-inducible splicing events in human pancreatic cancer organoids. **a.** Schematic of observed splicing event for EIF4A2, SRSF6 and FASTKD1. Arrows indicate qRT-PCR and qPCR primer locations **b.** Quantitative PCR comparing alternative isoform abundances for EIF4A2, SRSF6 and FASTKD1 in normoxia and hypoxia in PaCa-4, -6, -8 organoid lines. Arrows indicate isoform location. qPCR amplicons run on a 2% agarose gel. **c.** Isoform-specific qRT-PCR in PaCa-4, -6, -8 organoid lines (n = 3, shown is mean +/- SEM; *p < 0.05; **p < 0.01, ***p < 0.001, two-tailed unpaired t-test, n.s … not significant)

In the first round of validation, we could experimentally validate 3 HIF-dependent splicing events in eukaryotic initiation factor 4 A2 (EIF4A2), serine-argine rich splice factor 6 (SRSF6) and FAST kinase domain containing protein 1 (FASTKD1) out of 8 events tested (Supplementary Table T1) Alternative exon inclusion in EIF4A2 and SRSF6 are predicted to create non-coding transcripts, whereas FASTKD1 exon exclusion preserves the reading frame (Fig 3a). Quantitative PCR using primers located in the flanking exons of the alternative used exon for each event are visualized on a 2% agarose gel (Fig. 3b). Relative isoform abundance measured using boundary-spanning qRT-PCR primers are shown in (Fig. 3c).

However, in both TCGA cancer patients and our organoid system, the alternative splicing PSI changes observed were small and likely of limited biological importance. There was also inconsistency and variance in isoform abundances between the cell lines.

#### Coupled analysis HIF-dependent gene expression and alternative mRNA processing

While we were able to identify and validate differential splice events independent of changes in expression, we observed them to have a low effect size. Since HIF-induced transcriptional adaptation to hypoxia encompasses gene expression changes in thousands of genes, we decided to expand our initial analysis to identify events with significant changes in both splicing and expression (Supplementary Figure S2). In this expanded set, we identified more (654) events (Supplementary figure 2a) as well as overall larger effect sizes between conditions.

After another round of validation experiments, we experimentally validated HIF-dependent isoform changes for FAM13A, SLC35A3, ANKZF1, ANKDR37, CIDEB and NSMCE4A out of 26 events tested (Supplementary Table T1).

Next, we decided to focus our efforts on further characterizing the hypoxia-induced alternative splicing of SLC35A3, an octahelical transmembrane Golgi transporter for UDP-N-acetylglucosamine. SLC35A3 was among the top-ranked candidates after SplAdder analysis, identified in the TCGA patient cohort and was significantly associated with the hypoxia score in at least one cancer type, showed consistent hypoxia-dependent splicing in qRT-PCR validation in several model systems, and its biological role has been underexplored in the scientific literature.

We measured a strong HIF-dependent upregulation of SCL35A3 transcripts containing exon1-exon2 junctions (bs1-2) when compared to alt-exon2 (alt) (Fig. 4b). To estimate total isoform abundances, we performed quantitative PCR using alternative exon flanking primers (Fig. 4c, top). We observed a marked increase in skipping of the alternative and exon located between exon 1 and exon 2 of SLC35A3 upon hypoxia in PaCa-8 and PaCa-6, but not PaCa-4, as visualized on a 2% agarose gel (Fig. 4c, bottom). However, we noted that PaCa-4 already showed a strong baseline exon skipping and there was a discernible reduction in transcript abundance of alternative-exon containing longer isoforms in hypoxia when compared to normoxia. For all cell lines, we observed an exon-specific abundance increase of exon 1 during hypoxia, which is likely the result of an alternative transcription start site (TSS). During normoxia, transcription of SLC35A3 preferentially starts at a noncoding exon upstream of exon 1, whereas during hypoxia, transcription preferentially starts at exon 1. Mechanistically, the usage of an alternative TSS site coupled with an increase in exon skipping during HIF activation results in an overexpression of exon-1 containing long isoform of SLC35A3 (SLC35A3-L) over the canonical shorter SLC35A3 isoform. A schematic of the proposed isoform switch during HIF activation is shown in Figure 4d.

**Figure 4.**
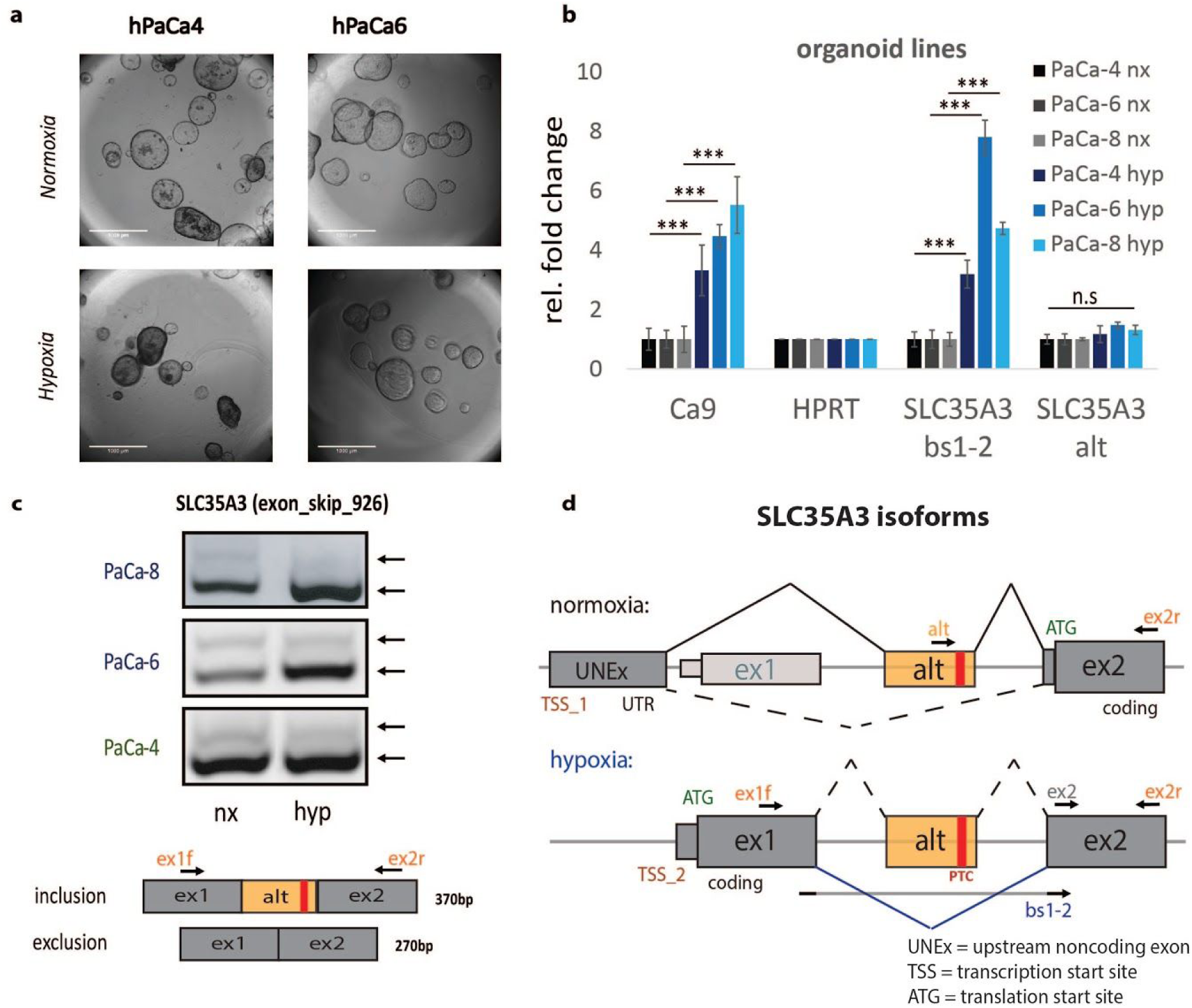
Alternative mRNA processing of SLC35A3 isoforms during hypoxia. **a**, Human cancer patient derived pancreatic organoids form spherical structures in matrigel in normoxia and after 24h 1% O_2_ hypoxia. **b**, SLC35A3 isoform-specific qRT-PCR in PaCa-4, -6, -8 organoid lines (n = 3, shown is mean +/- SEM; *p < 0.05; **p < 0.01, ***p < 0.001, two-tailed unpaired t-test, n.s … not significant) **c**, Quantitative PCR comparing SLC35A3 alternative exon inclusion in normoxia and hypoxia. Arrows indicate qPCR primer location. qPCR amplicons run on a 2% agarose gel. **d**, Schematic of hypoxia mediated SLC35A3 isoform switch and subsequent alternative mRNA processing. Arrows indicate primer locations.

### Hypoxia inducible splicing of SLC35A3 is dependent on both HIF1 and HIF2

We set out to identify if the observed alternative splicing of SLC35A3 is strictly dependent on HIF1 or HIF2 or if other cellular stresses related to hypoxia can affect their splicing pattern.

We treated the human PDAC lines AsPC-1 and PANC-1 for 24 and 48 hours (Supplementary Fig S4) with 1 mM deferoxamine (DFO), an iron chelator reported to induce pseudo-hypoxia by stabilizing HIF. As expected, we observe the transcriptional upregulation of HIF target genes CAIX, GLUT1 and APOL1 after DFO treatment (Supplementary Fig S3b) due to HIF stabilisation. For SLC35A3, both AsPC-1 and PANC-1 showed a significant upregulation of the boundary-spanning exon 1-2 junction (SLC35A3 bs1-2), while transcripts harboring the alternative exon (SLC35A3 alt) remained unchanged (AsPC-1) or even reduced (PANC1) under hypoxia, suggesting the observed isoform changes are not limited to one cell line.

To test time dependency, we set up AsPC-1 cells with and without ARNT knockdown and harvested RNA after 4, 24 and 48 hours of hypoxia (Fig 4a). Knockdown of ARNT with siRNA remained stable for all conditions (Fig 4a). GLUT1 is a known early HIF target gene and shows a slight transcriptional upregulation already after 4h and remaining constantly upregulated after 24h and 48h hours. We observed a strong induction of SLC35A3 alt-exon exclusion after 24h and 48h, but not after 4 hours of hypoxia, in accordance with our expectation of a HIF-dependent gene.

We wanted to find out if SLC35A3 splicing is dependent on HIF1α, HIF2α, or both. Therefore, we transduced AsPC-1 cells with a lentiviral construct carrying a short hairpin RNA against HIF1α (sh_HIF1a) or non-targeting control (sh_ns) and treated the cells to 1mM DFO for 24h (Fig 5b). We observe a significant reduction in HIF-target genes Ca9 and GLUT1 in sh_HIF1a cells compared to sh_ns. Correspondingly, SLC35A3 splicing during hypoxia was impaired, reducing usage of the SLC35A3 bs1-2 junction (exon exclusion) to baseline levels. Similarly, HIF2α knockdown in hypoxia reduced usage SLC35A3 bs1-2 junction, albeit not to baseline (Fig 5c).

**Figure 5.**
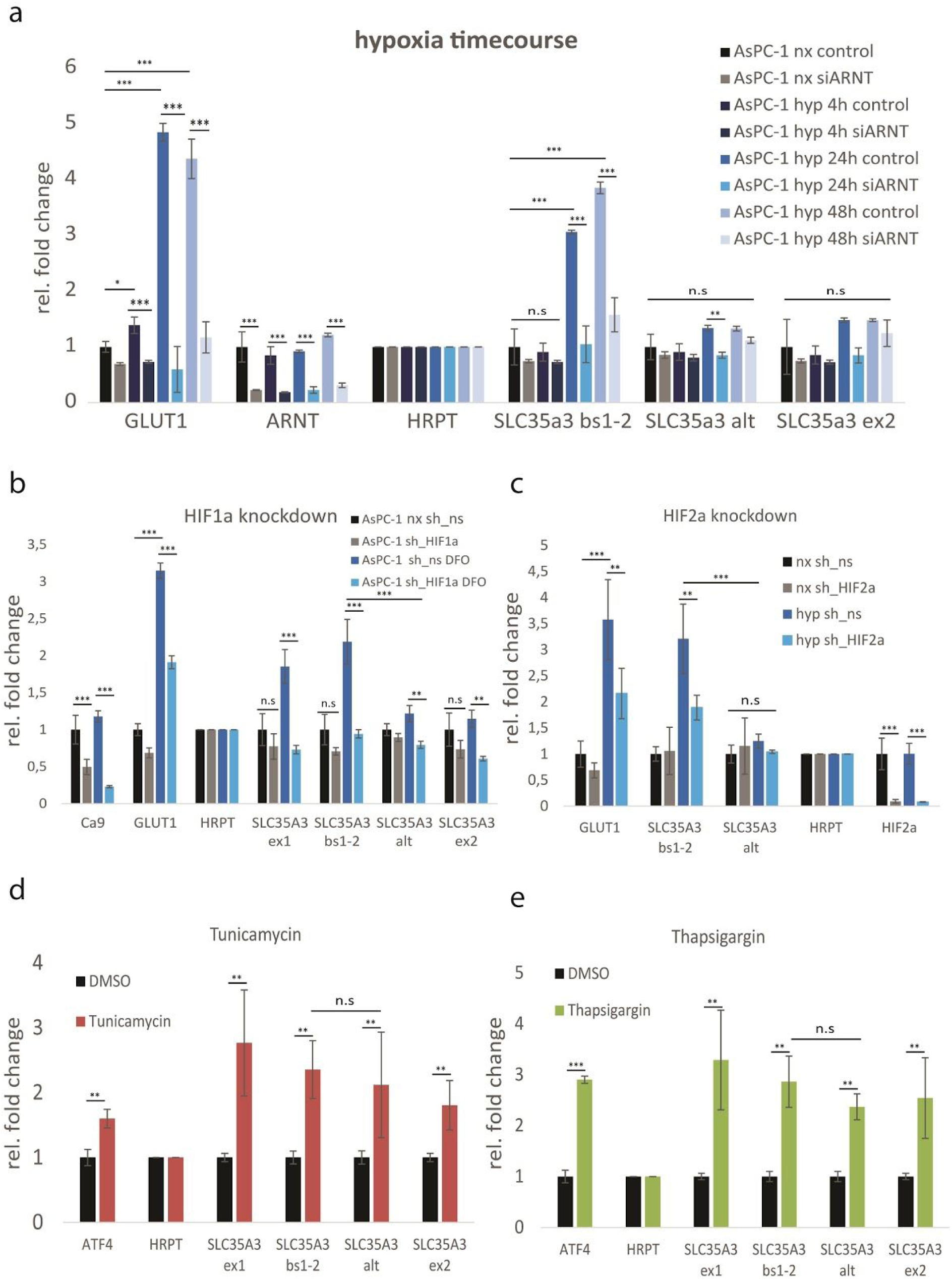
SLC35A3 dependency on hypoxia-inducible factors. **a.** Hypoxia timecourse. AsPC-1 cells were exposed 4, 24 or 48h to 1% O_2_ before harvest and SLC35A3 splicing assessment. (n = 3, shown is mean +/- SEM; *p < 0.05; **p < 0.01, ***p < 0.001, two-tailed unpaired t-test, n.s … not significant). **b & c**. SLC35A3 alternative exon usage comparing non-targeting shRNA control vs. shRNA mediated knockdown of HIF1α (b) or HIF2α (c) in hypoxia. HIF-dependent GLUT1 expressed is used for comparison. (n = 3, shown is mean +/- SEM; *p < 0.05; **p < 0.01, ***p < 0.001, two-tailed unpaired t-test, n.s … not significant) **d & e**. SLC35A3 alternative splicing in HUH7 cells treated with Tunicamycin (d) and Thapsigargin (e). (n = 3, shown is mean +/- SEM; *p < 0.05; **p < 0.01, ***p < 0.001, two-tailed unpaired t-test, n.s … not significant). Nx, normoxia; Hyp, hypoxia; siARNT; siRNA mediated knockdown of aryl-receptor nuclear translocator (ARNT); siControl, siRNA control (nontargeting)

Next, we investigated if other cellular stressors independent of HIF, could cause SLC35A3 splicing. SLC35A3 is a UDP-N-acetylglucosamine transporter located at the Golgi membrane [62]. The Golgi membrane is a sensitive organelle which harbors several stress kinases involved in the integrated stress response. Additionally, SLC35A3 upregulation has been reported in response to osmotic stress in CHO cells [63]. We reasoned that since both hypoxia and DFO are known to induce the integrated stress response, it is possible that the observed SLC35A3 splicing is a post-transcriptional response to cellular stress signaling. Therefore, we harvested RNA from HUH7 cells treated with Tunicamycin, which blocks N-linked glycosylation and induces the unfolded-protein response (UPR) (Fig 5d), and Thapsigargin, an inhibitor of sarco/endoplasmatic reticulum Ca^2+^ ATPase (SERCA) which causes ER stress (Fig 5e).

Surprisingly, both stress-inducing compounds cause an enrichment in total SLC35A3 mRNA, arguing for a role of this protein in the integrated stress response, but show no differences when comparing SLC35A3 isoforms including or excluding the hypoxia-dependent alternative exon.

We conclude that the observed hypoxia-inducible switch in SLC35A3 isoforms is strictly dependent on the presence of HIF proteins, but not other cellular stressors.

### Metabolic profiling of hypoxia-inducible SLC35A3-long isoform

SLC35A3 is a UDP-N-acetylglucosamine octahelical transmembrane transporter located at the Golgi membrane, potentially regulating flux through the hexosamine pathway. Therefore, we analyzed the mRNA expression of hexosamine pathway components, including the gate-keeping enzymes Glutamine-fructose-6-phosphate aminotransferase 1 & 2 (GFPT1 and GFPT2) responsible for converting glycolytic fructose-6-phosphate to D-glucosamine-6-phosphate using glutamine. We observed a marked increase in GFPT2 mRNA expression, but not GFPT1, and no change in downstream UDP-N-acetylglucosamine converting enzymes MGAT4A and OGT during hypoxia in PDAC organoids (Supplementary Fig. S4a). Next, we overexpressed two cDNA construct encoding SLC35A3-GFP (canonical isoform 1) and SLC35A3-L-GFP (isoform 3) in HUH7 cells assess where they would localize in cells (Supplementary Fig. S5). Both canonical and long isoform accumulated outside the nuclear membrane and had strong overlap with wheat germ agglutinin (WGA), a lectin with high affinity to N-acetylglucosamine often used as a Golgi marker.

To further detangle the role of hypoxia-dependent upregulation of SLC35A3 exon1-containing long isoforms (SLC35A3-L) we designed an siRNA directed at exon 1 of SLC35A3-L (Fig S4b), thereby allowing us to selectively deplete full-length transcripts (Fig S4c). Unexpectedly, we discovered that SLC35A3-L knockdown increased GFPT2 mRNA expression in both normoxia and hypoxia (Fig. 6d), but had no detrimental effect on the expression of other hexosamine pathway genes. GFPT2 mRNA expression levels have been correlated with UDP-N-acetylglucosamine levels previously [64], so we hypothesized that SLC35A3-L knockdown might influence steady state levels of this metabolite.

**Figure 6.**
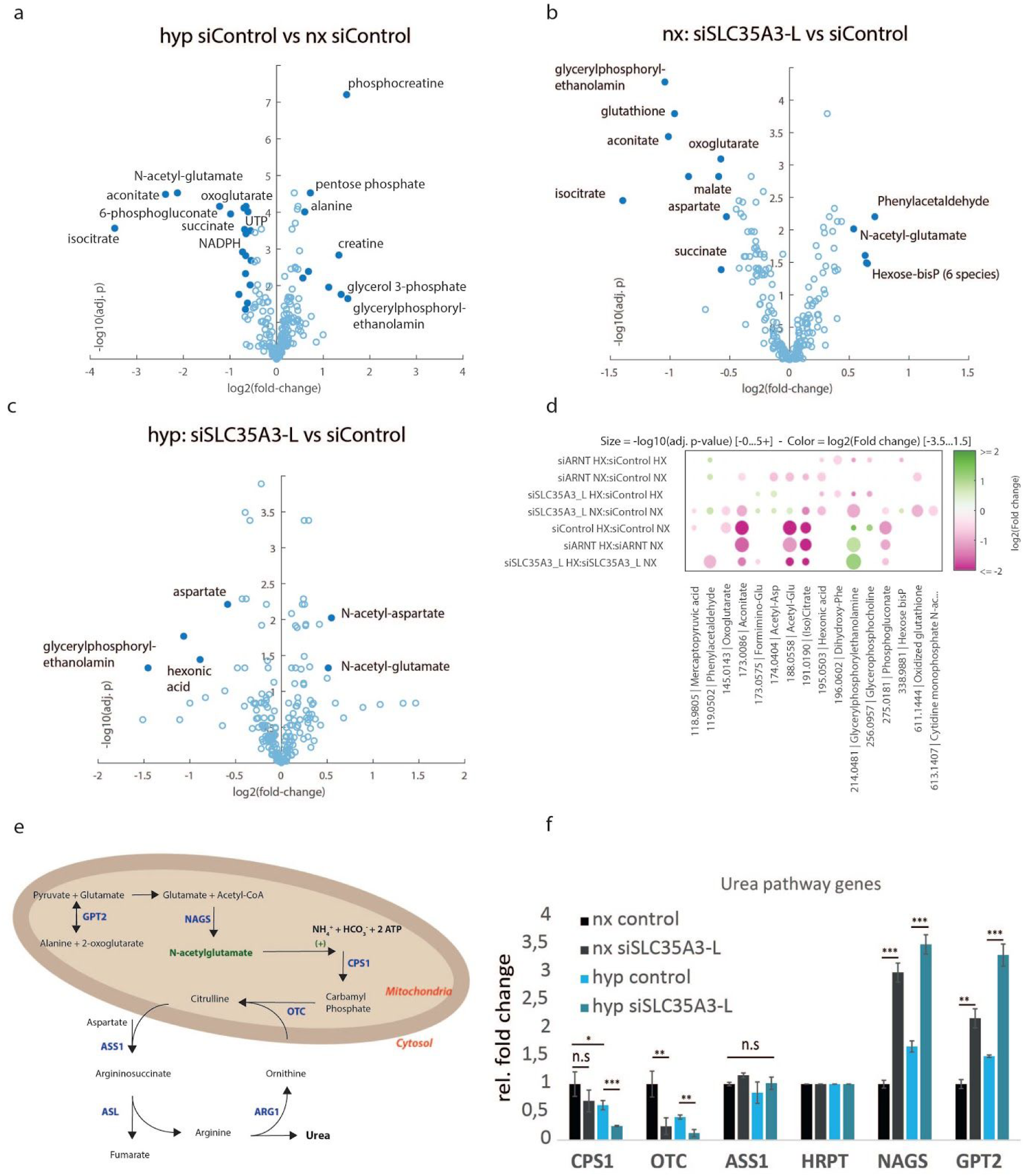
Metabolic profiling of HUH7 cells after SLC35A3-long knockdown. **a-c**. Volcano plots of pairwise comparison of metabolite abundances over experimental conditions in HUH7 cell with and without siRNA mediated knockdown of SLC35A3 isoform 3 exposed to (a) normoxia or (b) 24 hours of hypoxia. **d.** Summary of significantly changed metabolites over experimental conditions. **e.** Schematic of N-acetylglutamate pathway and urea cycle. **f.** qRT-PCR of urea pathway enzymes mRNA expression (n = 3, shown is mean +/- SEM; *p < 0.05; **p < 0.01, ***p < 0.001, two-tailed unpaired t-test, n.s … not significant). Nx, normoxia; Hyp, hypoxia; siSLC35A3L; siRNA mediated knockdown of the long isoform of SLC35A3; siControl, siRNA control (nontargeting)

Next, we assessed if SLC35A3-L knockdown has an impact on glycolysis or oxygen consumption using a SeaHorse XF24 Flux Analyzer. We transfected HUH7 and U2OS cells with siSLC35A3L or non-targeting siControl and cultured them with 1mM DFO or in normal DMEM medium (see Methods).

We found no changes in extracellular acidification (ECAR) or oxygen consumption (OCR) after siSLC35A3-L transfection in normoxia and hypoxia, indicating that SLC35A3-L knockdown is not impacting glycolytic flux or mitochondrial oxygen consumption rate (FigS4e-h).

To get a clearer picture on metabolite abundances, we performed metabolic profiling of HUH7 cells transfected with siControl, siARNT, and siSLC35A3-L and exposed these cells to normoxia or hypoxia for 24 hours. Hypoxia causes a consistent depletion of TCA cycle intermediates, such as (iso)citrate, succinate, malate, oxoglutarate as well as high energy metabolites like ATP, UTP or NADPH (Fig. 6a and supplementary table T2). Knockdown of SLC35A3-L during normoxia somewhat mimicked the depletion of TCA cycle intermediates like isocitrate, aconitate, succinate, malate, with the notable strong depletion of glutathione, a marker of oxidative stress. Interestingly, siRNA mediated knockdown of SLC35A3-L causes an accumulation (adj. p-value < 0.05) of N-acetyl-glutamate, rather than a depletion as during hypoxia. This accumulation is also significant when comparing isoform specific knockdown of SLC35A3-L to hypoxia control, while the knockdown had an overall minor effect on global metabolite homeostasis in this setting (Fig. 6c). Significant metabolite changes over all pairwise comparisons are summarized in Figure 6d.

Given the accumulation of N-acetyl-glutamate, we assessed the mRNA expression of N-acetyl-glutamate synthase (NAGS) and other enzymes related to the Urea pathway in HUH7 cells after isoform specific knockdown of SLC35A3L (Fig. 6e). We find upstream GPT2 and NAGS mRNA expression increased, partly explaining our accumulation of N-acetyl-glutamate (Fig. 6f). Furthermore, we observe a reduction in mRNA expression of downstream carbamyl phosphate synthase 1 (CPS1) and ornithine transcarbamylase (OTC), the gatekeeping enzymes to the Urea cycle pathway, following SLC35A3L knockdown or exposure to hypoxic conditions. Notably, the reduction in mRNA expression of these two enzymes is strongest when cells experience both hypoxia and SLC35A3-L knockdown (Fig. 6f).

## Discussion

Since the discovery of hypoxia-inducible factors two decades ago, hundreds of studies uncovered how this family of transcription factors influence a plethora of cellular processes through transcriptional upregulation of thousands of genes. Yet surprisingly little is known about the influence of HIF-proteins in post-transcriptional processes like pre-mRNA splicing.

In our HIF-perturbation experiment, we discovered a large set of hypoxia-inducible and HIF-dependent alternative splicing events using SplAdder. When we set out to experimentally confirm these findings and assess potential biological implications, we found that the relative abundance of the alternative spliced transcripts was in many cases just a fraction compared to the major isoforms. We cannot exclude that some of the discovered events play a biological role in response to hypoxia, however we reasoned that narrowing down our analysis to splicing events where identified isoforms make up an at least a 5% change in overall transcript abundance had a higher chance of having function impacts. To further strengthen our claim that candidate events were biologically or disease relevant, we sought to identify events that were not only observed in real patients in the TCGA cohort, but also correlated with the tumor’s hypoxia status. Establishing a tumor’s individual hypoxic score based on individual transcriptome data and associating HIF-dependent splicing events allowed us to identify and experimentally validate many promising candidate genes for follow up biological studies. To showcase the biological merits of this approach, we decided to study in-depth the HIF dependency of one promising candidate gene, SLC35A3.

For SLC35A3, we identified an alternative exon located in the very long intron between exon 1 and exon 2. The exon is conserved in various primate species, including chimpanzee, gorilla, orangutan, macaque and olive baboon, but in no other mammals. The canonical exon 1 and exon 2 are conserved in all mammalian species, indicating that this interspersed alternative exon is a rather recent addition. Interestingly, this alternative exon also harbors a premature termination codon, which would either lead to NMD-mediated RNA degradation of the transcript or cause the ribosome to stall and produce a truncated protein if translated. We also observe that SLC35A3 has a second start codon in exon 2, which produces a shorter SLC35A3 protein. According to both our transcriptome and available proteomic (https://www.proteomicsdb.org) data, this shorter isoform is the major isoform for SLC35A3, at least during normoxia. Under hypoxia, we observe a hypoxia-dependent increase in exon 1 usage and an overall enrichment of SLC35A3 transcripts including the splicing junction ex1-2, where the PTC-containing alternative exon is spliced out. This observation led us to speculate that alternative splicing might not be the only contributing factor to generate the observed splicing event, but that HIF proteins might be responsible to cause a shift in SLC35A3 transcription start site (TSS) usage, similar to what has been observed for the electron transport chain protein isoforms of COX4-1 and COX4-2 during hypoxia [65]. We observe in multiple model systems that HIF proteins play a crucial role in selecting for full-length exon-1 containing SLC35A3 transcripts which would otherwise be suppressed by transcription from an upstream TSS or by inclusion of an alternatively spliced-in PTC-containing exon.

At this point, it is unclear what function the addition of 42 N-terminal amino-acids encoded by exon 1 would serve. One plausible assumption is that these amino acids serve as a leader peptide, guiding SLC35A3-L towards the Golgi; however once we overexpressed both SLC35A3-L (containing exon1) and SLC35A3 (canonical isoform) we did not observe obvious differences in localization; both isoforms showed perinuclear and ER/Golgi association. SLC35A3 is known to heterodimerize with SLC35A2 (UDP-N-galactosamine transporter) [62], thus a leader peptide does not seem required for recruiting SLC35A3 towards the Golgi.

To address some of these questions, we selectively depleted SLC35A3-long isoforms during hypoxia and performed metabolic profiling and measured oxygen consumption and lactate production. Notably, we did not identify any changes in lactate production or oxygen consumption after selective downregulation of hypoxia-inducible SLC35A3 long isoform in both normoxia and when we stabilized HIF proteins with DFO, arguing for a neglectable impact of SLC35A3-long isoforms on influencing glycolysis or glycolytic flux.

Unexpectedly, we found an accumulation of mitochondrial metabolite N-acetyl-glutamate. NAG has been reported previously as a critical metabolite acting as catalytic cofactor for carbamyl phosphate synthase 1 (CPS1), the entry enzyme for conversion of ammonia to urea (urea cycle) in the liver [66,67]. Concomitantly, we also found a transcriptional upregulation of N-acetyl-glutamate synthase (NAGS) in both normoxia and hypoxia during SLC35A3L knockdown, as well as downregulation of CPS1 expression. The observed accumulation of NAG and the reduction in NAG-utilizing enzyme CPS1 as well as OTC suggest a decreased ability of SLC35A3-L depleted cells to get rid of excess nitrogen via the urea cycle. During hypoxia, it is beneficial for the cell to reduce urea cycle activity because urea production is an expensive ATP consuming process (CPS1 and ASS1 each use 2 ATP per catalytic reaction). A detailed role of SLC35A3-L in these metabolic processes is still unclear and will require further investigations.

One compelling hypothesis would attribute a hypoxia-protective role to HIF-induced SLC35A3-L, which might help cancer cells to deal with increased glutamine uptake during hypoxia mediated metabolic reprogramming and prevent nitrogen toxicity, a potential danger to cancer cells which rely on increased glutaminolysis [68].

In summary, our study shows how a genome wide splicing analysis from a perturbation sequencing experiment combined with pathway-stratified TCGA cohort data can be utilized to probe a transcription factor’s post-transcriptional influence. We rigorously validate a set of HIF-dependent splicing events in several model systems, including human pancreatic cancer patient derived organoids. Lastly, we demonstrate on the example of SLC35A3 how a HIF dependent mechanism can create an alternative isoform with implications in cancer metabolism.

## Supporting information

MP_diff_gene_expr_summary

MP_PCR_primer_list

MP_metabolic_data_curated

## AVAILABILITY

Gene expression data is available on Gene Expression Omnibus: https://www.ncbi.nlm.nih.gov/geo/query/acc.cgi?acc=GSE139673

Plotting and analysis scripts – https://github.com/ratschlab/hif_splicing_code_public

TCGA RNA-Seq and clinical data – https://portal.gdc.cancer.gov/

Spladder – https://github.com/ratschlab/spladder

DESeq2 – https://bioconductor.org/packages/release/bioc/html/DESeq2.html

ssPATHS – https://bioconductor.org/packages/devel/bioc/html/ssPATHS.html

MASS – https://cran.r-project.org/web/packages/MASS/index.html

topGO – https://bioconductor.org/packages/release/bioc/html/topGO.html

matchIt – https://cran.r-project.org/web/packages/MatchIt/index.html

## Acknowledgement

We would like to thank Dr. Andrea Aloia and Dr. Ilaria Guccini for reviewing the manuscript. We also would like to thank Dr. Werner Kovac for providing material and discussions for some cell culture experiments. We are grateful to Roger Meier and ScopeM for help with organoid microscopy. We also would like to express our gratitude towards Dr. Andre Kahles for generating and querying the splice graph for TCGA cohorts.

## Funding

This work was supported MSKCC core Funding to G.R.; ETH Zurich core funding to G.R and W.K.

## Conflict of interest

PM, ND, CKH, NZ, GS and GR declare no conflict of interest. CDC is a full time employee of Roche AG and a shareholder in Roche and AstraZeneca.

## Notes

#### Summary of Updates

Improved clarity of Fig3 and Fig4 Added citations

## References

1. Kaelin WG Jr, Ratcliffe PJ. Oxygen sensing by metazoans: the central role of the HIF hydroxylase pathway. Mol Cell. 2008;30: 393–402.

2. Samanta D, Prabhakar NR, Semenza GL. Systems biology of oxygen homeostasis. Wiley Interdisciplinary Reviews: Systems Biology and Medicine. 2017. p. e1382. doi:10.1002/wsbm.1382

3. Semenza GL. Oxygen sensing, homeostasis, and disease. N Engl J Med. 2011;365: 537–547.

4. Maltepe E, Schmidt JV, Baunoch D, Bradfield CA, Simon MC. Abnormal angiogenesis and responses to glucose and oxygen deprivation in mice lacking the protein ARNT. Nature. 1997;386: 403–407.

5. Arany Z, Huang LE, Eckner R, Bhattacharya S, Jiang C, Goldberg MA, et al. An essential role for p300/CBP in the cellular response to hypoxia. Proc Natl Acad Sci U S A. 1996;93: 12969–12973.

6. Neufeld G, Cohen T, Gengrinovitch S, Poltorak Z. Vascular endothelial growth factor (VEGF) and its receptors. FASEB J. 1999;13: 9–22.

7. Bunn HF, Poyton RO. Oxygen sensing and molecular adaptation to hypoxia. Physiol Rev. 1996;76: 839–885.

8. Goda N, Ryan HE, Khadivi B, McNulty W, Rickert RC, Johnson RS. Hypoxia-inducible factor 1alpha is essential for cell cycle arrest during hypoxia. Mol Cell Biol. 2003;23: 359–369.

9. Hackenbeck T, Knaup KX, Schietke R, Schödel J, Willam C, Wu X, et al. HIF-1 or HIF-2 induction is sufficient to achieve cell cycle arrest in NIH3T3 mouse fibroblasts independent from hypoxia. Cell Cycle. 2009;8: 1386–1395.

10. Kim J-W, Tchernyshyov I, Semenza GL, Dang CV. HIF-1-mediated expression of pyruvate dehydrogenase kinase: a metabolic switch required for cellular adaptation to hypoxia. Cell Metab. 2006;3: 177–185.

11. Wenger RH. Mammalian oxygen sensing, signalling and gene regulation. J Exp Biol. 2000;203: 1253–1263.

12. Papandreou I, Cairns RA, Fontana L, Lim AL, Denko NC. HIF-1 mediates adaptation to hypoxia by actively downregulating mitochondrial oxygen consumption. Cell Metab. 2006;3: 187–197.

13. Kothari S, Cizeau J, McMillan-Ward E, Israels SJ, Bailes M, Ens K, et al. BNIP3 plays a role in hypoxic cell death in human epithelial cells that is inhibited by growth factors EGF and IGF. Oncogene. 2003;22: 4734–4744.

14. Sermeus A, Michiels C. Reciprocal influence of the p53 and the hypoxic pathways. Cell Death Dis. 2011;2: e164.

15. Hubbi ME, Kshitiz, Gilkes DM, Rey S, Wong CC, Luo W, et al. A nontranscriptional role for HIF-1α as a direct inhibitor of DNA replication. Sci Signal. 2013;6: ra10.

16. Ryan HE, Lo J, Johnson RS. HIF-1 alpha is required for solid tumor formation and embryonic vascularization. EMBO J. 1998;17: 3005–3015.

17. Sowter HM, Ratcliffe PJ, Watson P, Greenberg AH, Harris AL. HIF-1-dependent regulation of hypoxic induction of the cell death factors BNIP3 and NIX in human tumors. Cancer Res. 2001;61: 6669–6673.

18. Semenza GL. HIF-1 mediates metabolic responses to intratumoral hypoxia and oncogenic mutations. J Clin Invest. 2013;123: 3664–3671.

19. Talks KL, Turley H, Gatter KC, Maxwell PH, Pugh CW, Ratcliffe PJ, et al. The expression and distribution of the hypoxia-inducible factors HIF-1alpha and HIF-2alpha in normal human tissues, cancers, and tumor-associated macrophages. Am J Pathol. 2000;157: 411–421.

20. Catalano V, Turdo A, Di Franco S, Dieli F, Todaro M, Stassi G. Tumor and its microenvironment: a synergistic interplay. Semin Cancer Biol. 2013;23: 522–532.

21. Philip B, Ito K, Moreno-Sánchez R, Ralph SJ. HIF expression and the role of hypoxic microenvironments within primary tumours as protective sites driving cancer stem cell renewal and metastatic progression. Carcinogenesis. 2013;34: 1699–1707.

22. Silva VL, Al-Jamal WT. Exploiting the cancer niche: Tumor-associated macrophages and hypoxia as promising synergistic targets for nano-based therapy. J Control Release. 2017;253: 82–96.

23. Parks SK, Cormerais Y, Pouysségur J. Hypoxia and cellular metabolism in tumour pathophysiology. J Physiol. 2017;595: 2439–2450.

24. Maxwell PH, Pugh CW, Ratcliffe PJ. Activation of the HIF pathway in cancer. Curr Opin Genet Dev. 2001;11: 293–299.

25. Brahimi-Horn C, Pouysségur J. The role of the hypoxia-inducible factor in tumor metabolism growth and invasion. Bull Cancer. 2006;93: E73–80.

26. Sena JA, Wang L, Heasley LE, Hu C-J. Hypoxia regulates alternative splicing of HIF and non-HIF target genes. Mol Cancer Res. 2014;12: 1233–1243.

27. Weigand JE, Boeckel J-N, Gellert P, Dimmeler S. Hypoxia-induced alternative splicing in endothelial cells. PLoS One. 2012;7: e42697.

28. David CJ, Manley JL. Alternative pre-mRNA splicing regulation in cancer: pathways and programs unhinged. Genes Dev. 2010;24: 2343–2364.

29. Zhang J, Manley JL. Misregulation of pre-mRNA alternative splicing in cancer. Cancer Discov. 2013;3: 1228–1237.

30. Kim E, Magen A, Ast G. Different levels of alternative splicing among eukaryotes. Nucleic Acids Res. 2007;35: 125–131.

31. Merkin J, Russell C, Chen P, Burge CB. Evolutionary dynamics of gene and isoform regulation in Mammalian tissues. Science. 2012;338: 1593–1599.

32. Cartegni L, Chew SL, Krainer AR. Listening to silence and understanding nonsense: exonic mutations that affect splicing. Nat Rev Genet. 2002;3: 285–298.

33. Wang G-S, Cooper TA. Splicing in disease: disruption of the splicing code and the decoding machinery. Nat Rev Genet. 2007;8: 749–761.

34. He C, Zhou F, Zuo Z, Cheng H, Zhou R. A global view of cancer-specific transcript variants by subtractive transcriptome-wide analysis. PLoS One. 2009;4: e4732.

35. Shkreta L, Bell B, Revil T, Venables JP, Prinos P, Elela SA, et al. Cancer-Associated Perturbations in Alternative Pre-messenger RNA Splicing. Cancer Treat Res. 2013;158: 41–94.

36. Chabot B, Shkreta L. Defective control of pre-messenger RNA splicing in human disease. J Cell Biol. 2016;212: 13–27.

37. Wang J, Dumartin L, Mafficini A, Ulug P, Sangaralingam A, Alamiry NA, et al. Splice variants as novel targets in pancreatic ductal adenocarcinoma. Sci Rep. 2017;7: 2980.

38. Kahles A, Ong CS, Zhong Y, Rätsch G. SplAdder: identification, quantification and testing of alternative splicing events from RNA-Seq data. Bioinformatics. 2016;32: 1840–1847.

39. Bolger AM, Lohse M, Usadel B. Trimmomatic: a flexible trimmer for Illumina sequence data. Bioinformatics. 2014;30: 2114–2120.

40. Dobin A, Davis CA, Schlesinger F, Drenkow J, Zaleski C, Jha S, et al. STAR: ultrafast universal RNA-seq aligner. Bioinformatics. 2013;29: 15–21.

41. Hartley SW, Mullikin JC. QoRTs: a comprehensive toolset for quality control and data processing of RNA-Seq experiments. BMC Bioinformatics. 2015;16: 224.

42. Love MI, Huber W, Anders S. Moderated estimation of fold change and dispersion for RNA-seq data with DESeq2. Genome Biol. 2014;15: 550.

43. Love MI, Anders S, Kim V, Huber W. RNA-Seq workflow: gene-level exploratory analysis and differential expression. F1000Res. 2015;4: 1070.

44. Benjamini Y, Hochberg Y. Multiple Hypotheses Testing with Weights. Scandinavian Journal of Statistics. 1997. pp. 407–418. doi:10.1111/1467-9469.00072

45. Ho DE, Imai K, King G, Stuart EA. MatchIt: Nonparametric Preprocessing for Parametric Causal Inference. Journal of Statistical Software. 2011. doi:10.18637/jss.v042.i08

46. Alexa A, Rahnenführer J, Lengauer T. Improved scoring of functional groups from gene expression data by decorrelating GO graph structure. Bioinformatics. 2006;22: 1600–1607.

47. Venables WN, Ripley BD. Modern Applied Statistics with S. Statistics and Computing. 2002. doi:10.1007/978-0-387-21706-2

48. Grossman RL, Heath AP, Ferretti V, Varmus HE, Lowy DR, Kibbe WA, et al. Toward a Shared Vision for Cancer Genomic Data. New England Journal of Medicine. 2016. pp. 1109–1112. doi:10.1056/nejmp1607591

49. ssPATHS (development version). In: Bioconductor [Internet]. [cited 30 Oct 2019]. Available: https://bioconductor.org/packages/devel/bioc/html/ssPATHS.html

50. Silver N, Cotroneo E, Proctor G, Osailan S, Paterson KL, Carpenter GH. Selection of housekeeping genes for gene expression studies in the adult rat submandibular gland under normal, inflamed, atrophic and regenerative states. BMC Mol Biol. 2008;9: 64.

51. Broutier L, Andersson-Rolf A, Hindley CJ, Boj SF, Clevers H, Koo B-K, et al. Culture and establishment of self-renewing human and mouse adult liver and pancreas 3D organoids and their genetic manipulation. Nat Protoc. 2016;11: 1724–1743.

52. Lin-Moshier Y, Marchant JS. A rapid Western blotting protocol for the Xenopus oocyte. Cold Spring Harb Protoc. 2013;2013. doi:10.1101/pdb.prot072793

53. Vorrink SU, Domann FE. Regulatory crosstalk and interference between the xenobiotic and hypoxia sensing pathways at the AhR-ARNT-HIF1α signaling node. Chem Biol Interact. 2014;218: 82–88.

54. Koshiji M, Kageyama Y, Pete EA, Horikawa I, Barrett JC, Huang LE. HIF-1alpha induces cell cycle arrest by functionally counteracting Myc. EMBO J. 2004;23: 1949–1956.

55. Ortmann B, Druker J, Rocha S. Cell cycle progression in response to oxygen levels. Cell Mol Life Sci. 2014;71: 3569–3582.

56. Chen C, Pore N, Behrooz A, Ismail-Beigi F, Maity A. Regulation of glut1 mRNA by hypoxia-inducible factor-1. Interaction between H-ras and hypoxia. J Biol Chem. 2001;276: 9519–9525.

57. Bowler E, Porazinski S, Uzor S, Thibault P, Durand M, Lapointe E, et al. Hypoxia leads to significant changes in alternative splicing and elevated expression of CLK splice factor kinases in PC3 prostate cancer cells. BMC Cancer. 2018;18: 355.

58. Venables WN, Ripley BD. Modern Applied Statistics with S-Plus. Springer; 2013.

59. Baker LA, Tiriac H, Clevers H, Tuveson DA. Modeling pancreatic cancer with organoids. Trends Cancer Res. 2016;2: 176–190.

60. Weeber F, Ooft SN, Dijkstra KK, Voest EE. Tumor Organoids as a Pre-clinical Cancer Model for Drug Discovery. Cell Chem Biol. 2017;24: 1092–1100.

61. Muthuswamy SK. Organoid Models of Cancer Explode with Possibilities. Cell stem cell. 2018. pp. 290–291.

62. Maszczak-Seneczko D, Sosicka P, Kaczmarek B, Majkowski M, Luzarowski M, Olczak T, et al. UDP-galactose (SLC35A2) and UDP-N-acetylglucosamine (SLC35A3) Transporters Form Glycosylation-related Complexes with Mannoside Acetylglucosaminyltransferases (Mgats). J Biol Chem. 2015;290: 15475–15486.

63. Lee JH, Jeong YR, Kim Y-G, Lee GM. Understanding of decreased sialylation of Fc-fusion protein in hyperosmotic recombinant Chinese hamster ovary cell culture: N-glycosylation gene expression and N-linked glycan antennary profile. Biotechnol Bioeng. 2017;114: 1721–1732.

64. Coomer M, Essop MF. Differential hexosamine biosynthetic pathway gene expression with type 2 diabetes. Mol Genet Metab Rep. 2014;1: 158–169.

65. Fukuda R, Zhang H, Kim J-W, Shimoda L, Dang CV, Semenza GL. HIF-1 regulates cytochrome oxidase subunits to optimize efficiency of respiration in hypoxic cells. Cell. 2007;129: 111–122.

66. Caldovic L, Ah Mew N, Shi D, Morizono H, Yudkoff M, Tuchman M. N-acetylglutamate synthase: structure, function and defects. Mol Genet Metab. 2010;100 Suppl 1: S13–9.

67. Shi D, Min L, Jin Z, Allewell NM, Tuchman M. Crystal structure of N-acetylglutamate synthase from Neisseria gonorrhoeae complexed with coenzyme A and L-glutamate. 2008. doi:10.2210/pdb3d2m/pdb

68. Yang L, Venneti S, Nagrath D. Glutaminolysis: A Hallmark of Cancer Metabolism. Annu Rev Biomed Eng. 2017;19: 163–194.

