## Supplementary material for "Characterisation of HIF-dependent alternative isoforms in pancreatic cancer"

#### Supplementary Figures

##### Figure S1

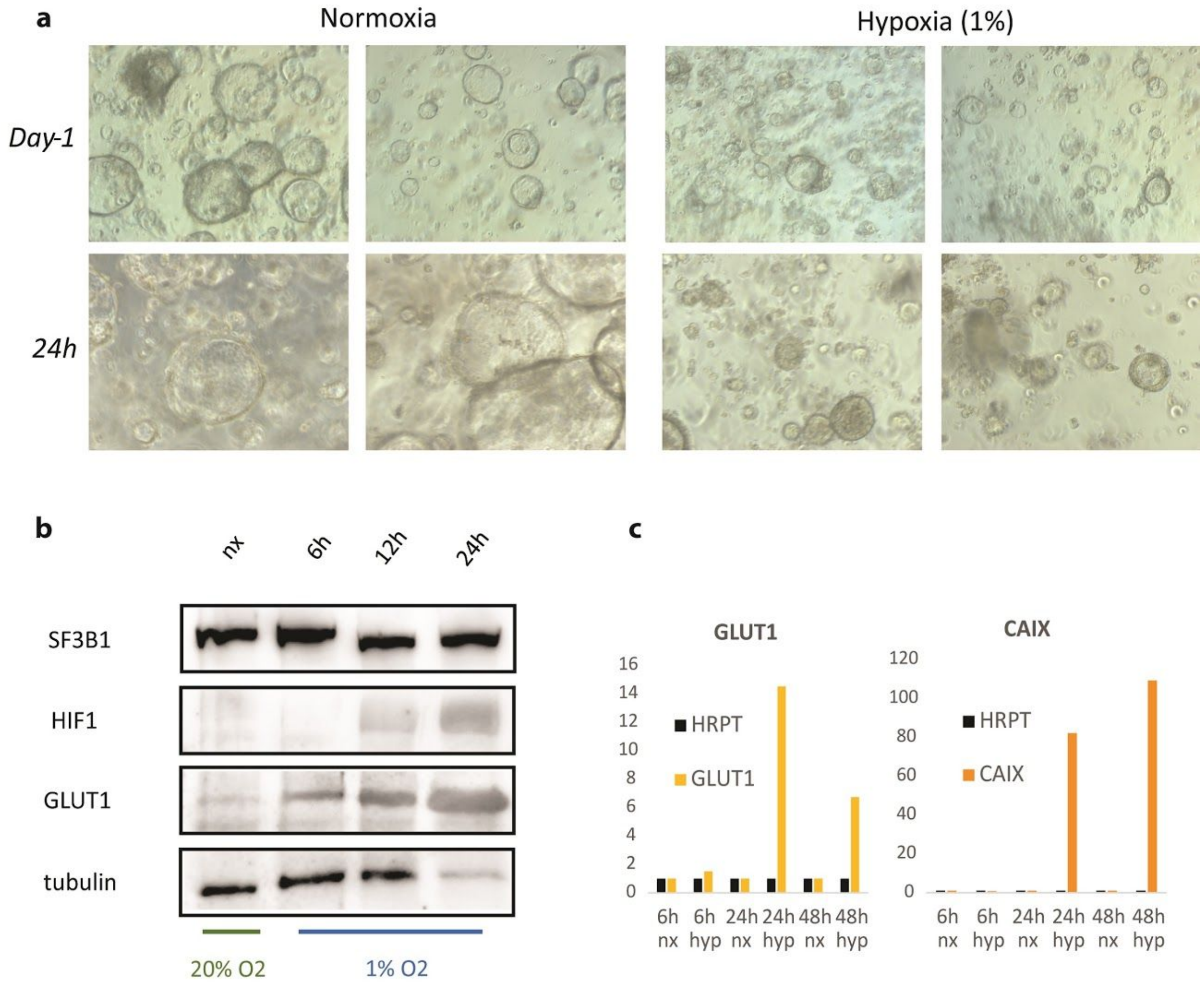

Supplementary Fig. S1**. PaCa-4 characterization. a.** Light microscopy of PaCa-4 organoid line in normoxia and hypoxia. **b**. Western blot of HIF1, GLUT1 and SF3B1 from PaCa-4 lysates after 0, 6, 12 and 24 hours of hypoxia. **c**. PaCa-4 qRT-PCR of canonical HIF target genes GLUT1, CAIX and APOL1 after 6, 24 and 48h of hypoxia. (n = 1 per timepoint)

### Figure S2

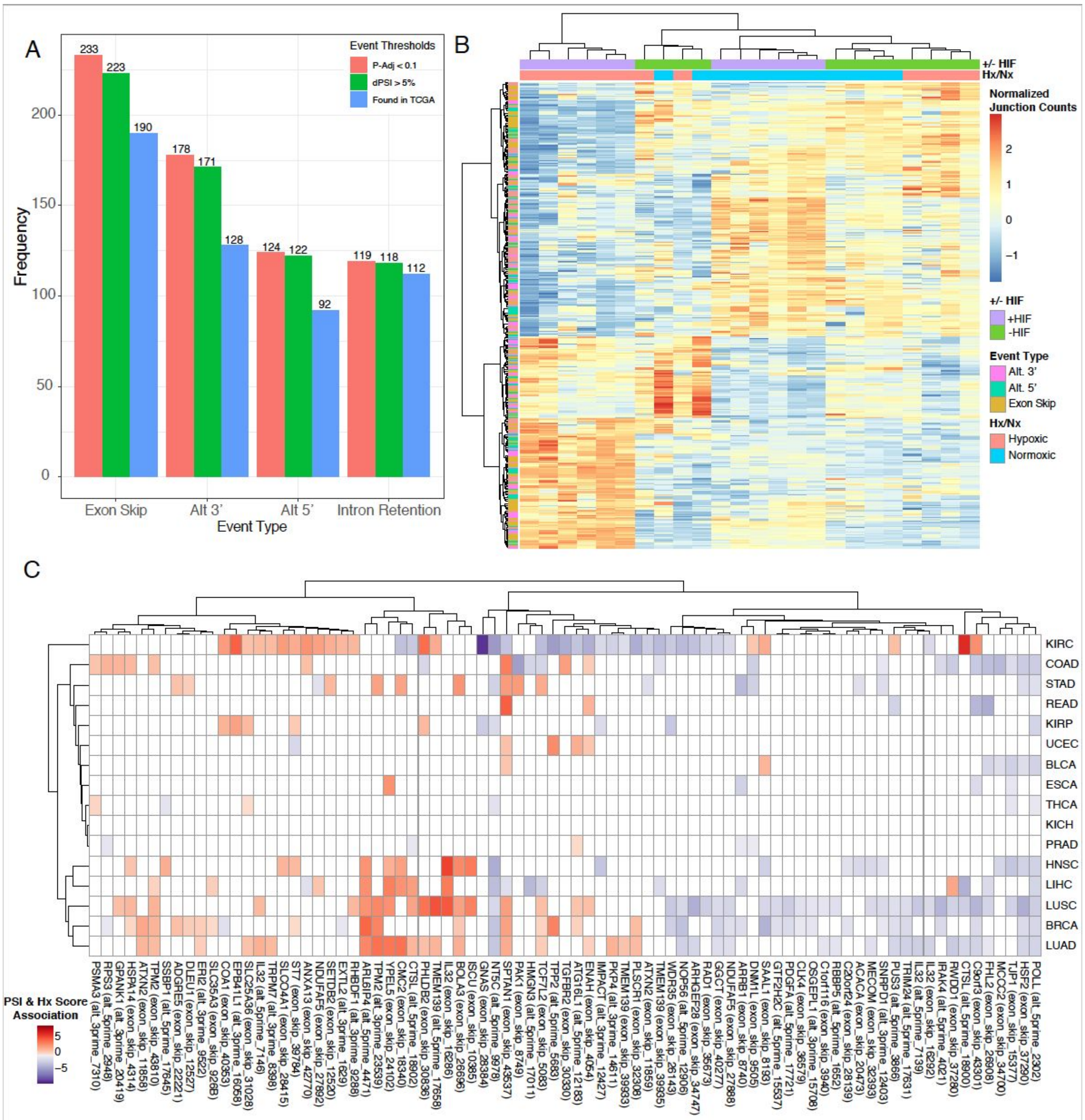

Supplementary Fig. S2. **Expression and splicing analysis**. **a.** Number of alternative splicing events identified using p-adjusted (pink) alone or p-adjusted and PSI >5% (green) combined as threshold. Number of remaining events after adding the additional filter that the events are observed in TCGA are depicted in blue. **b.** Heat map of centered and scaled splice junction and expression counts for identified alternative splicing events after hierarchical clustering. **c.** Heatmap of association results between PSI of our events of interest and an estimated hypoxia score in the TCGA cohort.

### Figure S3

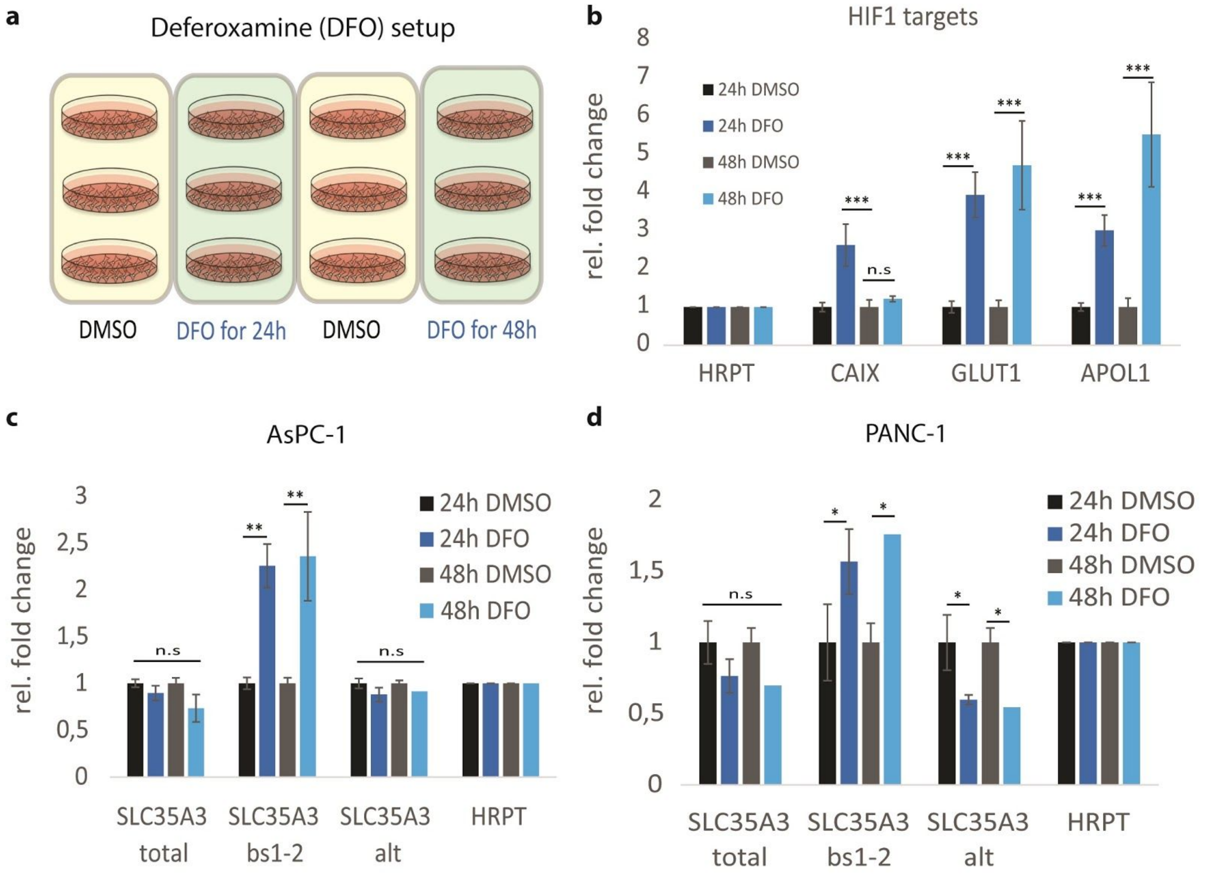

Supplementary Fig. S3**. Hypoxia-induced alternative splicing of SLC35A3**. **a,** Experimental setup. **b,** HIF1 target gene induction by DFO treatment. **c-d** SLC35A3 alternative splicing in AsPC-1 and PANC-1 PDAC cell lines treated with DFO for 24 and 48h. (n = 3, shown is mean +/- SEM; *p < 0.05; **p < 0.01, ***p < 0.001, two-tailed unpaired t-test, n.s … not significant)

### Figure S4

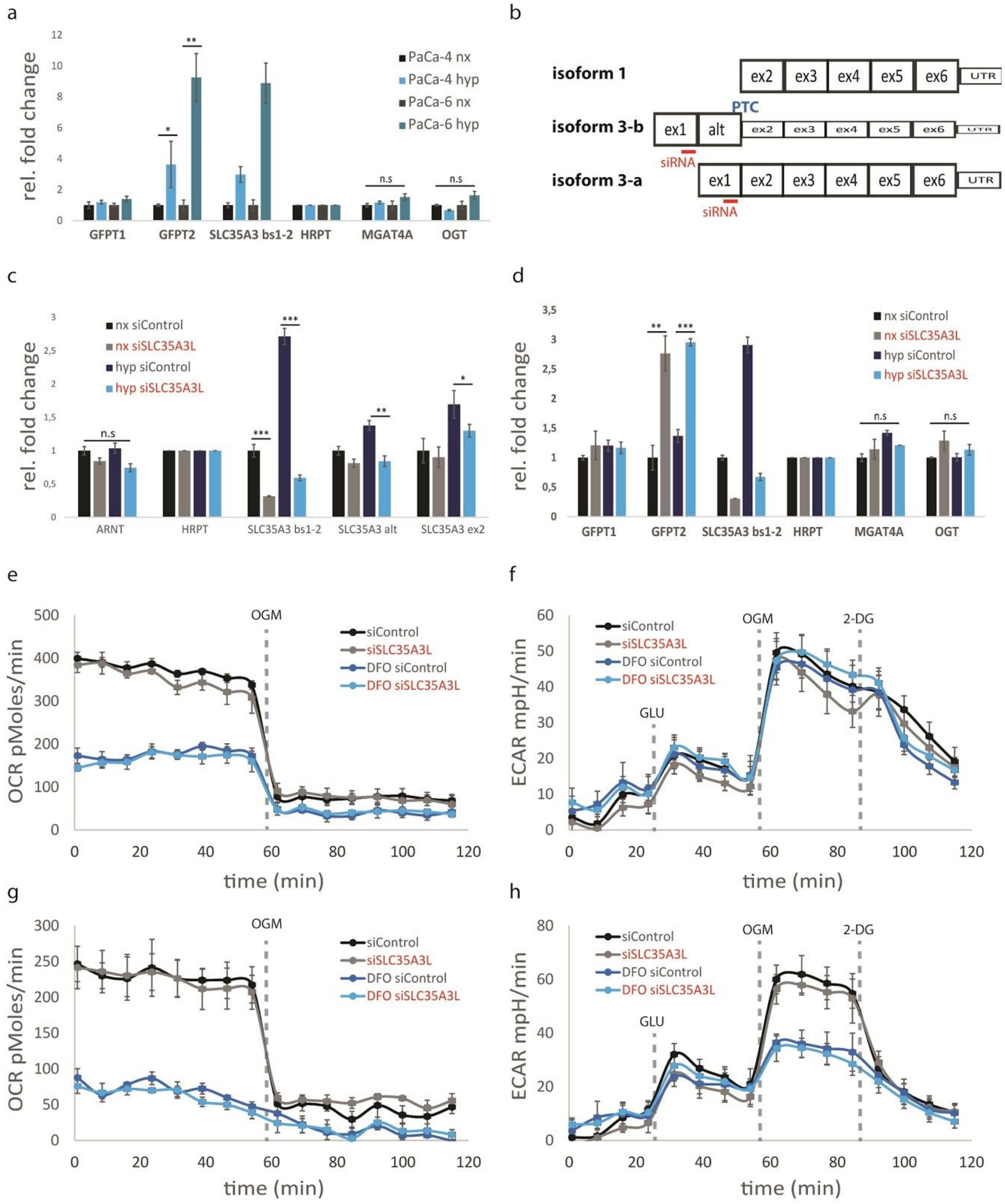

Supplementary Fig S4**. Selective knockdown of hypoxia-inducible SLC35A3-long isoform. a.** Assessment of glucosamine pathway genes in human PDAC organoids in normoxia and hypoxia. GFPT2 … Glutamine-fructose-6-phosphate aminotransferase 2. (n = 3, shown is mean +/- SEM; *p < 0.05; **p < 0.01, ***p < 0.001, two-tailed unpaired t-test, n.s … not significant) **b.** Schematic of protein-coding SLC35A3 isoform 1 and isoforms 3-a/b. siRNA position for selective knockdown are indicated in red. **c.** qRT-PCR quantification of siRNA targeted SLC35A3-exon1 containing isoforms. (n = 3, shown is mean +/- SEM; *p < 0.05; **p < 0.01, ***p < 0.001, two-tailed unpaired t-test, n.s … not significant) **d**. Assessment of hexosamine pathway genes after long-isoform specific SLC35A3 knockdown in HUH7 cells in normoxia and hypoxia. (n = 3, shown is mean +/- SEM; *p < 0.05; **p < 0.01, ***p < 0.001, two-tailed unpaired t-test, n.s … not significant). **e-h**. Measurement of oxygen consumption (e,g) and extracellular acidification (f,h) during glycolytic-stress test program for HUH7 (e,f) and U2OS (g,h) in normoxia or DFO-induced hypoxia with and without selective depletion of SLC35A3-long isoform. Spaced lines indicate time points of injection. GLU… glucose. OGM… oligomycin. 2-DG… 2-deoxyglucose.

### Figure S5

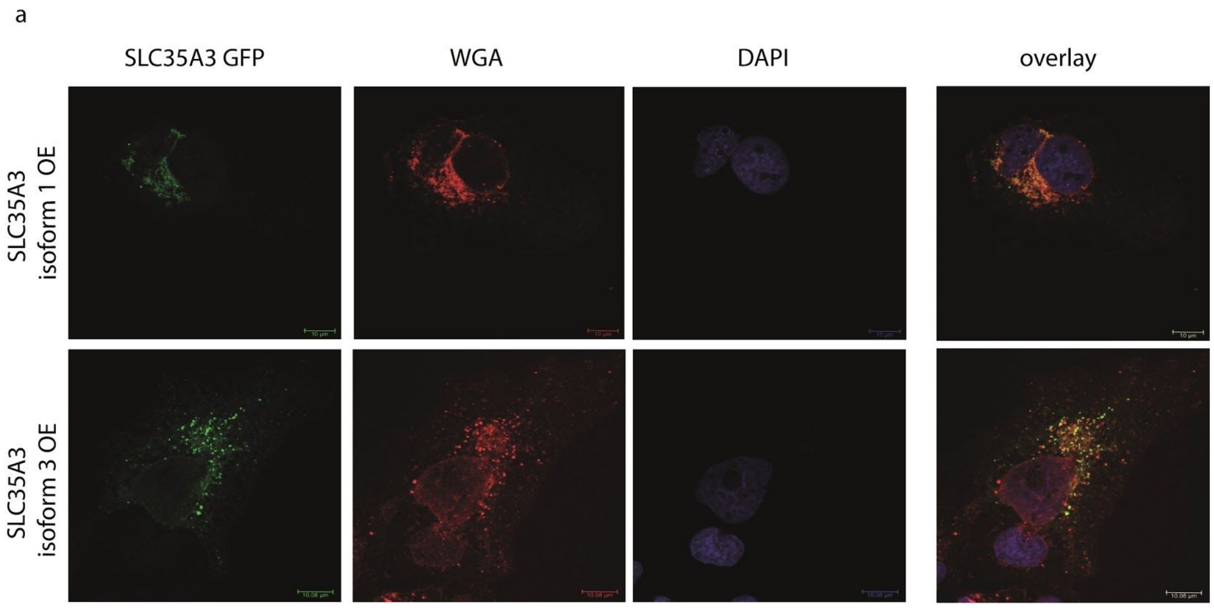

Supplementary Fig S5**. Localisation of SLC35A3 isoforms 1 and 3.** Overexpression of a cDNA construct encoding SLC35A3(isoform 1)-GFP (top) and SLC35A3(isoform 3)-GFP (bottom) in HUH7 cells. Fixated cells were stained with anti-GFP (green), wheat-germ agglutinin (WGA, red) and 4′,6-diamidino-2-phenylindole (DAPI, blue).

### Supplementary Tables

#### Table T1

| Validations summary | | | | |
|---|---|---|---|---|
| **Gene** | **qRT-PCR** | **alt. splicing** | **biol. relevance** | **consistent ov**[illegible] |
| Analysis I : splicing -expression effects | | | | |
| EIF4A2 | yes | yes | yes | no |
| AFF4 | yes | no | | |
| SRSF6 | yes | yes | yes | no |
| FUS | yes | no | | |
| FASTKD1 | yes | yes | yes | no |
| SRRM2 | yes | no | | |
| HNRNPH1 | yes | no | | |
| ANLN | yes | no | | |
| Analysis II : splicing + expression effects | | | | |
| P4HA1 | yes | no | | |
| AldoC | yes | yes | no | |
| PTPRR | yes | yes | no | |
| NPAS2 | yes | yes | no | |
| CTPS1 | yes | no | | |
| FAM13A | yes | yes | yes | yes |
| KDM1A | yes | no | | |
| LDHA | yes | no | | |
| MCM8 | yes | no | | |
| SLC35A3 | yes | yes | yes | yes |
| LRR1 | yes | no | | |
| ZSWIM8 | yes | yes | no | |
| DIMT1 | yes | yes | no | |
| HP1G2 | yes | no | | |
| ANKZF1 | yes | yes | yes | no |
| PTPRH | yes | yes | no | |
| CIDEB | yes | yes | yes | no |
| ANKRD37 | yes | yes | yes | no |
| NSMCE4A | yes | yes | yes | no |
| FADS3 | no | | | |
| NPSR1 | yes | no | | |
| CCDC107 | no | | | |
| RPS2 | no | | | |
| TUBAL3 | no | | | |
| RPS9 | no | | | |
| IL32 | no | | | |

**Supplementary Table 1**: Overview of Spladder candidate alternative splicing events chosen for qRT-PCR (column 2) and quantitative PCR (column 3) validation. Biological relevance (column 4) assessment was based on estimated likelihood/potential of alternative/divergent isoform function. Consistency (column 5) was called only when all tested cell systems showed similar isoform behavior.

### Supplementary Data

#### Gene omnibus

RNA sequencing data can be accessed via gene omnibus. For detailed information consult "MP_geneomnibus_submission.xlsx".

#### Differential gene expression results

The differential gene expression analysis results are collected in the summary spreadsheet "MP_diff_gene_expression_summary.xlsx" for each pairwise comparison between conditions.

#### Expression_Independent_Differential_Splicing_Results.zip

This zipped folder contains the results of the expression independent differential splicing analysis for each event type. The differential splicing results for each event are in the following files: exon_skip.tsv, alt_5prime.tsv, alt_3prime.tsv, and intron_retention.tsv. The remaining file, sig_event_positions_v2.tcga_PSI.tsv, contains the PSI values for the events of interest across all TCGA patients.

#### Expression_Dependent_Differential_Splicing_Results.zip

This zipped folder contains the results of the expression dependent differential splicing analysis for each event type. The differential splicing results for each event are in the following files: exon_skip.tsv, alt_5prime.tsv, alt_3prime.tsv, and intron_retention.tsv. The remaining file, sig_event_positions_v2.tcga_PSI.tsv, contains the PSI values for the events of interest across all TCGA patients.

#### PCR primer sequences

All qPCR and qRT-PCR primer sequences used in this study can be found in the file "MP_PCR_primer_list.xlsx".

#### Metabolic profiling

The file "MP_metabolomics_DATA CURATED.xlsx" contains log2fold metabolite changes with adjusted p-values for each pairwise comparison between conditions.
